# RIBO-former: leveraging ribosome profiling information to improve the detection of translated open reading frames

**DOI:** 10.1101/2023.06.20.545724

**Authors:** Jim Clauwaert, Zahra McVey, Ramneek Gupta, Gerben Menschaert

## Abstract

Ribosome profiling is a deep sequencing technique used to chart translation by means of mRNA ribosome occupancy. It has been instrumental in the detection of non-canonical coding sequences. Because of the complex nature of next-generation sequencing data, existing solutions that seek to identify translated open reading frames from the data are still not perfect. We propose RIBO-former, a new approach featuring several innovations for the *de novo* annotation of translated coding sequences. RIBO-former is built using recent transformer models that have achieved considerable advancements in the field of natural language processing. The presented deep learning approach allows to omit several pre-processing steps as features are automatically extracted from the data. We discuss various steps that improve the detection of coding sequences and show that read length information of all mapped reads can be leveraged to improve the predictive performance of the tool. Our results show RIBO-former to outperform previous methodologies. Additionally, through our study we find support for the existence of translated non-canonical ORFs, present along existing coding sequences or on long non-coding RNAs. Furthermore, several polycistronic mRNAs with multiple translated coding regions were detected.

## 1 Introduction

Ribosome profiling was first introduced by Ingolia et al. to study the translatome of cells through deep sequencing (1). The technique sequences ribosome-protected fragments that are aligned to a reference genome or transcriptome. The resulting profile quantifies the ribosome occupancy at nucleotide resolution, serving as a proxy for translation in the evaluated sample. A decade of ribosome profiling data has proven the tool to be a driving factor for the detection of coding sequences (CDSs) (2, 3, 4, 5, 6, 7, 8). It has successfully underlined the identification and workings of translational control, differential splicing (9), microproteins (10), and drug mechanisms (11, 12).

Similar to the output generated by other next-generation sequencing techniques, ribosome profiling results in complex data that involves various processing steps. Biological variation, usage of different translation inhibitors and lab protocols and sequencing depth are known variables that affect the fingerprint created by ribosome profiling experiments (13). Today, it is standard practice to re-align reads by their length when distilling features from the ribosome data (Supplementary Table A1). This practice has resulted from two important observations. First, the read length influences the A/P-site (within the ribosome complex) offset calculation of the ribosome-protected fragment (RPF) (1). Second, the count of RPFs (pinpointed to this offset position) within the coding sequence follows a strong triplet periodicity following the the reading frame of translated coding sequences (Supplementary Figure A4-7). Several approaches exist that determine custom offsets per read length in order to align the majority of reads to the same reading frame. This is achieved by evaluating the aligned reads using a set of well-characterized coding sequences (14, 15, 16). Tools applying ribosome profiling data still suffer from a high number of false positives. Other types of validation, such as mass spectrometry, are still widely applied to confirm the existence of newly proposed coding sequences (3, 4, 8).

Deep learning techniques are an important asset for handling large quantities of raw unprocessed data as their ability to automatically extract relevant features makes them extremely effective at mapping correlations between variables. These are, as such, well-suited to process next-generation sequencing data generated from a ribosome profiling experiment. We propose RIBO-former, a deep learning approach to detect translated coding sequences from ribosome profiling data (Figure 1). RIBO-former is designed to omit several pre-processing steps, including the creation of an ORF library, re-alignment of reads by read length, and feature generation. Through extensive evaluation, we find our approach to have the best predictive performance when including ribosome read length information alongside all aligned reads. RIBO-former offers several other advantages that can facilitate future scaling and application of the tool for ORF delineation. For example, the model is modular, annotates the full transcriptome, and can be pre-trained on a large selection of other ribosome profiling data, which was shown to improve performances. Our benchmark proves RIBO-former to outperform existing tools by a substantial margin. Furthermore, model predictions support the translation of non-canonical ORFs (ncORFs), such as present on long noncoding RNA (lncRNA), or upstream (overlapping) (u(o)ORF), downstream (overlapping) (d(o)ORF), and within the bounds (intORF) of known canonical coding sequences. Alongside, multiple occurrences of polycistronic mRNA are detected, yielding more than one translated open reading frame within the same transcript.

**Figure 1:**
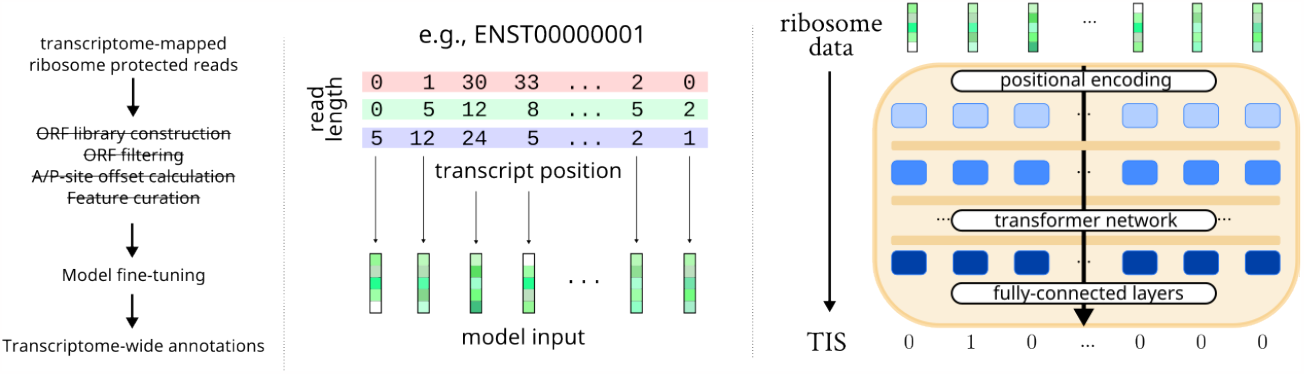
Overview of RIBO-former pipeline. (left) Different steps performed by RIBO-former. Steps that are typically performed by other tools but omitted by RIBO-former are striked through. (middle) RIBO-former processes ribosome reads along a transcript region and generates vector embeddings for each transcript position based on the number of mapped reads and read length fractions. (right) Illustration of the RIBO-former model. Expressed translation initiation sites (TISs) are predicted as a proxy of coding sequences. After generating the input vector embeddings, the data is sent through a transformer model. A set of fully-connected layers returns a probability score for each position on the transcript.

## 2 Material and Methods

RIBO-former is created to map the translatome at single-nucleotide resolution of samples using ribosome profiling data. To simplify the experimental set-up, we train a model to detect active translation initiation sites (TISs) from which the open reading frame can be derived. RIBO-former processes ribosome profiling data along a full transcript (Figure 1). No pre-processing of other types of data, such as start codon or ORF information, are used to curate features or build a candidate ORF library. Instead, all positions on the transcriptome are evaluated. RIBO-former is built upon previous work (TIS transformer), where a transformer model is optimized to detect translation initiation sites from transcript sequence data (17). Mapping of the ribosome profiling data is performed using STAR (18) and cutadapt (19) (Supplementary Files, Section 1.1).

### 2.1 Input data generation

Several approaches in parsing ribosome data have been evaluated. The first strategy uses ribosome read count information of reads mapped to a single location. In accordance with the standard practice (Supplementary Table A1) of processing ribosome data, custom offsets are determined for each read length. Read lengths ranging from 20 to 40 nucleotides are included in the data. To allow computation with a transformer-based architecture, vector representations that function as embeddings are used as inputs to the model. Vector embeddings map concepts within a multi-dimensional numerical space, where their positioning with respect to other embeddings (distance, angle) allow meaningful computations. Specifically, the vector embedding **e**_**c**_ for a given position is obtained from the read count *c* using a set of feed-forward layers *ϕ*.

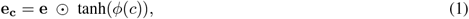

with *c* ∈ [0, 1], *ϕ* : ℝ^1^ *→* ℝ^*h*^, and **e** ∈ ℝ^*h*^. Read counts are first normalized across the transcript for numerical stability. The initial vector embedding **e** is optimized as part of the training process. *h* is a hyperparameter of the model indicating the input dimension.

A new strategy explores the inclusion of ribosome read length information as part of the information applied to determine TIS locations. No offsets are applied, and each read is mapped by its 5’-position. For a given transcript position, **e**_**l**_ is calculated using the read length fractions **l** following the equation:

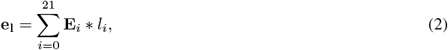

with **E** ∈ ℝ^21*×h*^ and **l** ∈ [0, 1]^21^, where Σ1 = 1. The matrix **E** incorporates vector embeddings for read lengths 20–40 and is optimized as part of the training process. Note that ribosome data by read length is sparse and the majority of values in **l** are 0.

In this study, we evaluate models trained using either **e**_**c**_ or **e**_**c**_ + **e**_**l**_ as input to the transformer model. Supplementary Figures A8–9 illustrate the generation of input vectors from ribosome profiling data for both strategies.

### 2.2 Model architecture and optimization

Continuing upon our previous work on detecting TISs using transcript sequence information (17), we utilize an identical model framework for RIBO-former. An important building block of this architecture is a recent innovation in calculating full attention introduced by Choromanski et al. (20), allowing long-range attention spanning the full transcripts. A detailed description of the network architecture is given in section 3.3 of the Supplementary Files. A binary cross-entropy loss is used to optimize the model with.

Selection of a final model architecture was achieved using a hyperparameter selection set-up (Supplementary Table A5). Hyperparameter tuning was performed using the ribosome data set featuring the most mapped reads (SRR2733100). No single parameter was observed to have a substantial impact on the performance. Nonetheless, the total number of model parameters was shown to correlate with the properties of the loss curve of the validation set (Supplementary Figure A10).

### 2.3 Data selection and evaluation

A myriad of ribosome profiling data sets were used covering a variety of tissues and treatment methods (cycloheximide, harringtonine, and lactimidomycine), including cells without antibiotic treatment (3, 4, 5, 6, 7, 8, 21, 22, 23, 24, 25, 26, 27, 28, 29) (Supplementary Table A2). All evaluated data are human. Supplementary Table A3 lists the number of reads mapped at different steps throughout the study, revealing the different data sets to cover a wide range of mapped reads.

The full transcriptome constitutes of 251,121 transcripts, with a total of 431,011,438 positions. The aim of this study is to predict the translatome, based on ribosome profiling information. In this study, Ensembl GRCh38 version 107 is used to positively label known TISs. Varying approaches exist that evaluate the number of mapped reads in order to reduce the positive set of the proteome to match that of a likely translatome. Filtering the positive set incurs an unnecessary factor of noise, as selection of thresholds, and thus inclusion of existing translation initiation sites into the positive set, is pseudo-arbitrary. Furthermore, sub-setting the input data or altering the labels can hinder the utility of the tool and future benchmarking efforts. We have decided to use all known translation initiation sites to set up a positive set. This does affect the maximum performance a tool can achieve using a certain data set, as an unknown amount of coding sequences are not being translated or have simply not been captured by the ribosome profiling experiment. Nonetheless, it does not hinder comparison between different tools. A higher performance score indicates one approach to have reconstructed a larger part of the translatome using ribosome profiling data, supporting its superiority over the other.

## 3 Results

### 3.1 Read length information can be leveraged to improve performances

While the length of mapped reads serves as an important factor determining the positioning of the RPF within the ribosome complex (14), the application of read length offsets fails to capture the full complexity of RPF alignments. This is apparent as a large quantity of reads does not adhere to one selected reading frame (Supplementary Figure A7), and variability exists between profiles of different read lengths (Supplementary Figure A4–6). Furthermore, no previous study has disproven the existence of a more complex correlation between biological factors and the behavior of ribosome read lengths along a transcript.

For all eight data sets, we have trained RIBO-former using ribosome profiling data with and without ribosome length information (see Material and Methods). Except for the few hundred weights used to compute the input vector, all approaches apply the same model architecture and thus number of model parameters (*∼*220K). In addition to utilizing the 5’ position to map reads, Plastid (14) and RiboWaltz (15) offsets were evaluated due to their widespread adoption and continued code maintenance.

Results show a consistent increase in performance for models incorporating ribosome read length information (Figure 2, Supplementary Table A6, Supplementary Figures A11-A13). Surprisingly, when only applying read count information, optimal performances are achieved when reads are not offset by their read length (i.e., 5’ positions). Otherwise, RiboWaltz shows to perform better than Plastid. When incorporating read length information, a larger number of correlations can be learned between the input data when a sufficient number of reads are present. The difference between performances for different data sets reflects these additional data requirements, where a higher read depth (Supplementary Figure A3) of the ribosome data results in larger jumps in performance when including read length information.

**Figure 2:**
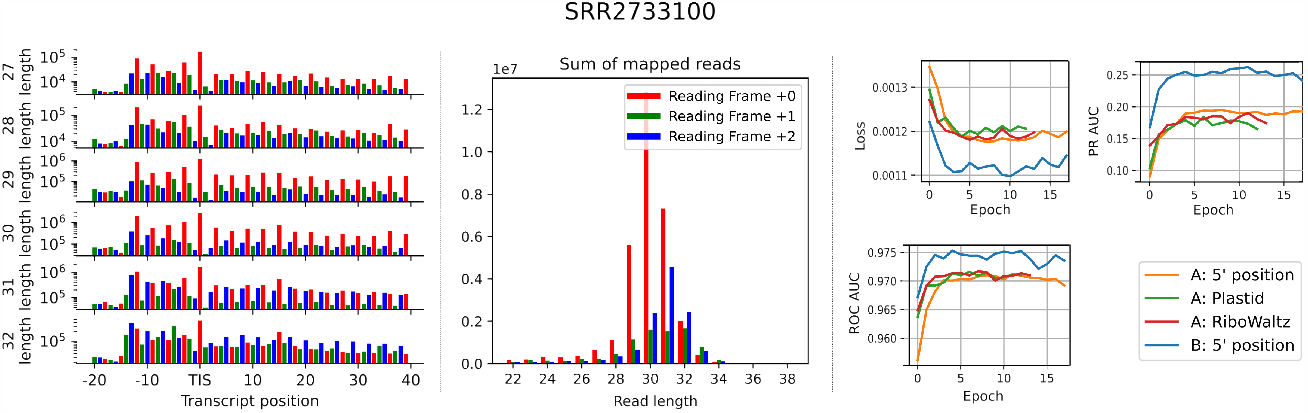
Read length information can be leveraged to improve detection of translated coding sequences. Data is from SRR2733100. (left) Counts of reads mapped by their 5’ positions along translation initiation sites listed in the consensus coding sequence library (CCDS). These showcase unique patterns of read alignments per read length. Read counts are taken by only evaluating translation initiation sites of coding sequences within the consensus coding sequence (CCDS) library. (middle) Read length counts binned by reading frame. This highlights the skewed abundance of reads as influenced by the reading frame of the neighboring translation initiation site. Patterns are unique for each data set and read length. (right) Validation loss, area under the receiver operating characteristic curve (ROC AUC), and area under the precision recall curve (PR AUC) per epoch for different input token strategies throughout the training process. Results indicate the relevance of read length information for the prediction of translation initiation sites using ribosome profiling data, especially for data sets featuring a higher read depth (see Supplementary Figure A3). Strategy A generates input tokens for the model utilizing only read count information for every position of the transcript. Reads are offset by read length following three strategies (5’, Plastid, and RiboWaltz). Strategy B includes information on both the positions and read lengths of the mapped reads.

### 3.2 Pre-training RIBO-former improves performance and training times

Ribosome profiling data is influenced by various factors, such as lab protocols, treatments, sample input material (cell types and/or species) and sequencing depth. RIBO-former, being a deep learning approach, relies on leveraging complex interactions between ribosome profiling patterns found within data. It does therefore not perform well on data it is not trained on, as learned correlations do not generalize well between data sets. To achieve high predictive performance of the tool, RIBO-former has to be trained on parts of the data set it needs to evaluate. However, to perform *de novo* detection of translated CDSs, a machine learning model should not be applied on regions of the transcriptome it is trained on. Thus, at least two models need to be optimized on non-overlapping folds of the transcriptome to cover it fully.

In order to make RIBO-former pertinent for future use, we evaluated a model that was first trained on a plethora of ribosome profiling data sets. Pre-trained models have been successfully used to improve performances and training times. By pre-training the model with a combination of several ribosome profiling data sets, it is able to discover general RIBO-seq correlations that are shared between different data sets. A selection of eight complementary ribosome profiling experiments were selected to pre-train the model. These cover samples obtained from various different samples treated with different translation inhibitors (Supplementary Table A2).

Two types of pre-trained models have been evaluated. The first set of pre-trained models is optimized to detect TISs, following an identical learning objective as outlined above. The second set of models is optimized using a self-supervised learning setting, where labels are derived from the input data. This is a popular approach in natural language processing, producing models capable of handling a variety of tasks, In this setting, random locations of the input are masked, where the model is tasked to impute the presence of ribosome reads at these positions (Supplementary Figure A14).

The results show that, compared to training models from scratch, improved performances and drastically reduced training times are achieved when further training (i.e., fine-tuning) the pre-trained model on our previously evaluated data sets (Supplementary Table A7, Supplementary Figures A15–A16). Models pre-trained using the supervised learning setting were shown to be the most successful. Importantly, models achieved convergence after only one iteration of the data (also referred to as training epoch), meaning optimization of the model can be achieved within one hour on modern hardware using a single graphical processing unit.

### 3.3 RIBO-former outperforms existing tools

Several tools exist that utilize ribosome profiling data in various manners to delineate translated ORFs (Supplementary Table A1). Tools like RiboTaper(8), riboHMM (30), and Scikit-ribo (31) are packages that combine both RNA-seq and Ribo-seq data to achieve this goal. Other software, such as PROTEOFORMER (32) and RiboTISH (33), can leverage information from the combination of cycloheximide and lactimidomycin or harringtonine treated data sets.

In this study, we benchmark RIBO-former with PRICE (34), Rp-Bp (35), RiboCode (16), RiboTISH (33), and Ribotricer (36). These tools were selected based on various factors: previously reported benchmarks, the presence of continued support for a tool through software updates and code maintenance, and their adoption by the community (16, 35, 36). Unlike RIBO-former, existing methods are designed to only evaluate a subset of transcriptome positions. As RIBO-former provides predictions along the full transcriptome, it is possible to provide a one-by-one comparison on the full subset of evaluated positions provided by the other tools. Post-processing steps that further filter the output predictions of all tools have been omitted, as these are not related to the predictive capability of the models. For example, this includes steps that filter down the results to only include the longest possible ORF on a transcript featuring an ATG start codon. A full list with the commands used for each tool are listed in the Supplementary Files.

Using coding sequences annotated by Ensembl as the positive set, we show RIBO-former to outperform all other tools on all evaluated data sets (Figure 3, Supplementary Table A9, Supplementary Figure A17). RIBO-former especially contrasts itself from existing approaches when it is evaluated on large sets featuring few positive samples (Rp-Bp, RiboTISH, Ribotricer). Note that it is not correct to compare the performances between existing tools (e.g., RiboCode> Rp-Bp), as different sets of evaluated ORFs decide the complexity of the benchmark and, therefore, the resulting performance metric.

**Figure 3:**
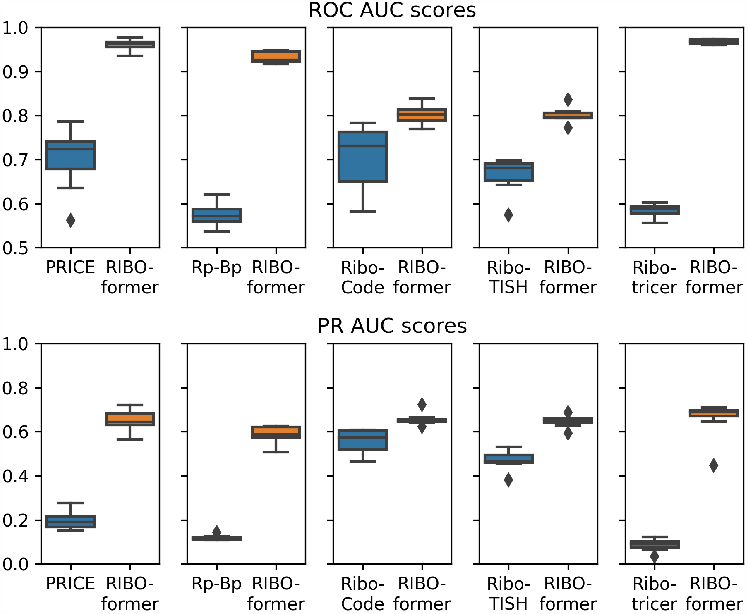
RIBO-former performances for *de novo* detection of expressed coding sequences as compared to existing tools. Given are the box plots using the performance on the eight evaluated data sets. Previous tools only evaluate a small selection of ORFs based on a variety of filters (e.g., start codon, read count). A one-by-one comparison for those positions is possible, as RIBO-former annotates the full transcriptome. With coding sequenes annotated by Ensembl as the positive set, the area under the receiver operating characteristic curve (ROC AUC) and area under the precision-recall curve (PR AUC) are calculated. Note that the number of positions and composition of positive and negative samples evaluated by each tool is unique and influences the score metric. Supplementary Table A9 gives a completer and more accurate overview of the benchmark listed by tool and data set, including metadata on the number of positions evaluated and size of the positive set.

### 3.4 Combination of replicates can improve read depth and aid detection

RIBO-former can successfully combine multiple runs of a ribosome profiling experiment to improve the detection of translated ORFs (Supplementary Table A3). These results follow the previous observation where our tool performs best for data sets featuring the highest read depth. Practically, reads are combined from multiple data sets through summation. Technical replicates are the most suitable for this approach, where no variability is expected between the translatome of multiple data sets and ribosome read profile characteristics are supposed to be similar. While biological replicates are bound to have more variations in the expression profiles of the the samples, correlations between mapped reads of the data sets are likely to be shared due to similarities in sample prep and other factors.

### 3.5 Evaluation of the non-canonical model predictions

Lastly, an evaluation is made of the top scoring model predictions with a focus on ncORFs (Figure 4). We observe that the model output distribution closely reflects the model uncertainty for each data set (Figure 4A). To illustrate, for a given threshold (e.g., 0.6), the precision of the positive set on all data sets is relatively similar (e.g., 0.5–0.6, blue bars in Figure 4A). As the PR/ROC AUC are both rank-based metrics, this property is not a given from our previous results. We furthermore observe a similar composition of ncORFs between the positive sets of the different data sets (orange bars in Figure 4A). For all data sets, u(o)ORFs are substantially more abundant than d(o)ORFs (green/red versus purple bars in Figure 4A).

**Figure 4:**
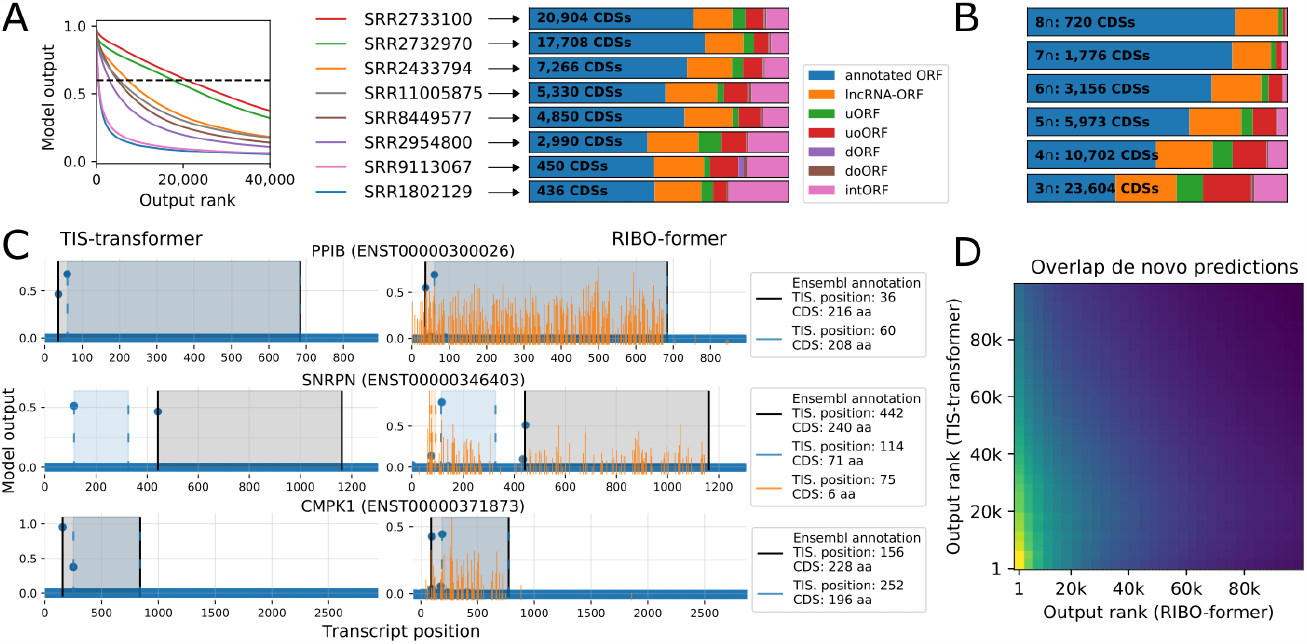
Analysis of the model outputs focusing on non-canonical predictions. (**A**) Model output as a function of output rank reflects a degree of model certainty captured by the model predictions. These are correlated to read depth of the data sets. For model predictions scoring above 0.6 (dotted line), the composition of open reading frame (ORF) types for each data set and total number of predicted coding sequences (CDSs) is given. (**B**) Taking a look at the predicted coding sequences that are present within top 100k (0.023%) predictions of multiple data set, the number and composition of coding sequences is given for instances shared between three (3∩) or more data sets. (**C**) Transcripts with multiple translation initiation sites are detected by both RIBO-former and TIS transformer (17). TIS transformer predicts TISs solely using transcript sequence information. Given are the model outputs (y-axis) for each position of the transcript (x-axis) for TIS transformer (left) and RIBO-former (rigth; data set SRR2732970). The bounds of the predicted coding sequence are given for high scoring sites. Mapped read counts used as input to RIBO-former are given in bright orange and displayed on a logarithmic scale. (**D**) Heatmap reflecting the fraction of intersecting predictions (higher: yellow) between different bins (per 2000) along the top 100k predictions of each tools. The figure reflects that higher scoring coding sequences of either tool are more likely to be shared within the top 100k predictions of the other tool. Note that the top scoring predictions for RIBO-former are obtained by merging the results of the eight data sets and removing lower-scoring duplicates.

The intersection of predictions between multiple ribosome experiments are another powerful approach to reduce the chance of false positives when selecting ncORFs (Figure 4B). To illustrate, utilizing the overlap between the top 100k predictions on each data set, a total of 128 ncRNAs have been detected on all data sets. The likelihood of eight models sharing a predicted CDSs within the top 100k predictions through random chance is only ∼ 10^−56^, as the top 100k predictions only constitute 0.023% of all predicted sites.

In a previous study, we introduced TIS transformer, a transformer-based neural network following the same design principles as RIBO-former. Both RIBO-former and TIS transformer are optimized to detect TIS sites at nucleotide-resolution level, albeit using different types of input data. TIS transformer only processes transcript sequence information where RIBO-former only processes ribosome profiling sequencing data. As such, both tools can be used to provide independent validation for the existence of alternative translation productions (for example from ncRNAs). We observe that similar prediction profiles exist along transcripts, including those featuring multiple high-scoring coding sequences (Figure 4C). This correlation exists on a macro level, where high-scoring predictions are more likely to be shared by both tools. Specifically, comparing the top 100k predictions of both tools, we identify that higher scoring predictions of one tool have a larger overlap with higher-scoring predictions of the other tool (Figure 4D).

## 4 Discussion

RIBO-former detects translation initiation sites from ribosome profiling read information along the transcript. Our model achieves this by providing predictions at every position of the transcriptome. We deem our approach to hold several advantages over previous tools. (i) The tool relies solely on the arrangement of ribosome-protected fragments (RPFs) to detect TISs. Additional information, such as sequence information (e.g., start codon, stop codon) or information on the properties of the open reading frame (e.g., length, number of reads mapped) could perpetuate potential biases within the decision-making process of the model. Filtering based on these properties can be achieved as a post-processing step, rather than a pre-processing step. (ii) The tool omits pre-processing steps that re-map reads as a function of their read length, pinpointing them to one position after an offset calculation. Based on analysis of the data, we know this step constitutes a loss of information or the introduction of biases. Leaving out this step results in higher model performances. These results are according to expectations as deep learning approaches are able to do automated feature extraction. (iii) The mapping of the full transcriptome allows for modular post-processing steps where specific sites of interest are guaranteed to be evaluated. Filters based on number of reads per transcript, sequence information, and ORF properties can be applied on the full set in line with the objectives of the user (Supplementary Table A8). Furthermore, evaluating the full transcriptome facilitates future benchmarking. (iv) The application of state-of-the-art machine learning models and the incorporation of ribosome length information outperforms all previously designed tools for all evaluated data sets. Importantly, the tool was shown to work well with a variety of data, covering a wide range of sequencing depths, sample types, and antibiotic treatments applied. We find these observations to support the utility of the tool. Although a focus was set on the human genome, RIBO-former can be applied across different species, and future findings might even prove the inclusion of data sets from multiple species to be a valid strategy to create a pre-trained model.

We deem the main disadvantage of RIBO-former to be the computational requirements to run the tool. Its reliance on a graphical processing unit is likely to be an important limiting factor for wide-spread adoption. To alleviate this requirement, we have investigated the use of pre-trained models which have substantially decreased training times (Supplementary Figure A15–16). The accessibility of cloud computing solutions can further provide support for the application of graphical processing unit-powered algorithms in labs that are otherwise lacking support of local hardware.

Future work will focus on calibrating the multiple models trained alongside one another and introducing post-processing steps. We believe the latter to be an important asset to further filter sites of interest and correct model inaccuracies. For example, as RIBO-former relies solely on ribosome profiling information, pinpointing translation initiation sites can suffer from low accuracy for transcripts featuring a lower number of mapped reads. Data shows model predictions can be off several nucleotides when compared to those annotated by Ensembl, a phenomenon especially prevalent for data sets featuring a lower read depth (Supplementary Figure A18). Post-processing steps can detect and correct likely inaccuracies by evaluating model scores and neighboring codon prevalence. Other post-processing strategies can introduce filters based on read count and ORF properties (Supplementary Table A8).

In this study, we propose RIBO-former as a new tool to delineate translated ORFs and find it to provide substantial improvements compared to existing approaches. Findings corroborate the translation from so-called non-coding transcripts (ncORFs) and the presence of multiple translated ORFs from one single transcript (polycistronic nature). Together with TIS transformer, we have now designed and provided two independent but complementary systems for the *de novo* delineation of coding sequences. We believe these tools to be well-positioned to spearhead the discovery or to provide validation on non-canonical coding sequences.

## 5 Data availability

All the data, scripts and model outputs are available for public use on GitHub (https://github.com/jdcla/RIBO_former_paper) and Zenodo (https://doi.org/10.5281/zenodo.8059446). Resources and instructions required to apply the RIBO-former framework and train models on new data are also available on GitHub https://github.com/jdcla/RIBO_former. A python package is created to support this work (https://pypi.org/project/transcript-transformer/).

## References

1. Ingolia, N. T., Ghaemmaghami, S., Newman, J. R. S., and Weissman, J. S. (April, 2009) Genome-Wide Analysis in Vivo of Translation with Nucleotide Resolution Using Ribosome Profiling. Science, 324(5924), 218–223.

2. van Heesch, S., Witte, F., Schneider-Lunitz, V., Schulz, J. F., Adami, E., Faber, A. B., Kirchner, M., Maatz, H., Blachut, S., Sandmann, C.-L., Kanda, M., Worth, C. L., Schafer, S., Calviello, L., Merriott, R., Patone, G., Hummel, O., Wyler, E., Obermayer, B., Mücke, M. B., Lindberg, E. L., Trnka, F., Memczak, S., Schilling, M., Felkin, L. E., Barton, P. J. R., Quaife, N. M., Vanezis, K., Diecke, S., Mukai, M., Mah, N., Oh, S.-J., Kurtz, A., Schramm, C., Schwinge, D., Sebode, M., Harakalova, M., Asselbergs, F. W., Vink, A., de Weger, R. A., Viswanathan, S., Widjaja, A. A., Gärtner-Rommel, A., Milting, H., dos Remedios, C., Knosalla, C., Mertins, P., Landthaler, M., Vingron, M., Linke, W. A., Seidman, J. G., Seidman, C. E., Rajewsky, N., Ohler, U., Cook, S. A., and Hubner, N. (June, 2019) The Translational Landscape of the Human Heart. Cell, 178(1), 242–260.e29.

3. Ji, Z., Song, R., Regev, A., and Struhl, K. (December, 2015) Many lncRNAs, 5’UTRs, and Pseudogenes Are Translated and Some Are Likely to Express Functional Proteins. eLife, 4, e08890.

4. Raj, A., Wang, S. H., Shim, H., Harpak, A., Li, Y. I., Engelmann, B., Stephens, M., Gilad, Y., and Pritchard, J. K. (May, 2016) Thousands of Novel Translated Open Reading Frames in Humans Inferred by Ribosome Footprint Profiling. eLife, 5, e13328.

5. Martinez, T. F., Chu, Q., Donaldson, C., Tan, D., Shokhirev, M. N., and Saghatelian, A. (April, 2020) Accurate Annotation of Human Protein-Coding Small Open Reading Frames. Nature Chemical Biology, 16(4), 458–468.

6. Gaertner, B., van Heesch, S., Schneider-Lunitz, V., Schulz, J. F., Witte, F., Blachut, S., Nguyen, S., Wong, R., Matta, I., Hübner, N., and Sander, M. (August, 2020) A Human ESC-based Screen Identifies a Role for the Translated lncRNA LINC00261 in Pancreatic Endocrine Differentiation. eLife, 9, e58659.

7. Chen, J., Brunner, A.-D., Cogan, J. Z., Nuñez, J. K., Fields, A. P., Adamson, B., Itzhak, D. N., Li, J. Y., Mann, M., Leonetti, M. D., and Weissman, J. S. (March, 2020) Pervasive Functional Translation of Noncanonical Human Open Reading Frames. Science, 367(6482), 1140–1146.

8. Calviello, L., Mukherjee, N., Wyler, E., Zauber, H., Hirsekorn, A., Selbach, M., Landthaler, M., Obermayer, B., and Ohler, U. (February, 2016) Detecting Actively Translated Open Reading Frames in Ribosome Profiling Data. Nature Methods, 13(2), 165–170.

9. Reixachs-Solé, M., Ruiz-Orera, J., Albà, M. M., and Eyras, E. (April, 2020) Ribosome Profiling at Isoform Level Reveals Evolutionary Conserved Impacts of Differential Splicing on the Proteome. Nature Communications, 11(1), 1768.

10. Weaver, J., Mohammad, F., Buskirk, A. R., and Storz, G. (March, 2019) Identifying Small Proteins by Ribosome Profiling with Stalled Initiation Complexes. mBio, 10(2), e02819–18.

11. Chu, J. and Pelletier, J. (June, 2018) Therapeutic Opportunities in Eukaryotic Translation. Cold Spring Harbor Perspectives in Biology, 10(6), a032995.

12. Wolfe, A. L., Singh, K., Zhong, Y., Drewe, P., Rajasekhar, V. K., Sanghvi, V. R., Mavrakis, K. J., Jiang, M., Roderick, J. E., Van der Meulen, J., Schatz, J. H., Rodrigo, C. M., Zhao, C., Rondou, P., de Stanchina, E., Teruya-Feldstein, J., Kelliher, M. A., Speleman, F., Porco, J. A., Pelletier, J., Rätsch, G., and Wendel, H.-G. (September, 2014) RNA G-quadruplexes Cause eIF4A-dependent Oncogene Translation in Cancer. Nature, 513(7516), 65–70.

13. Mudge, J. M., Ruiz-Orera, J., Prensner, J. R., Brunet, M. A., Calvet, F., Jungreis, I., Gonzalez, J. M., Magrane, M., Martinez, T. F., Schulz, J. F., Yang, Y. T., Albà, M. M., Aspden, J. L., Baranov, P. V., Bazzini, A. A., Bruford, E., Martin, M. J., Calviello, L., Carvunis, A.-R., Chen, J., Couso, J. P., Deutsch, E. W., Flicek, P., Frankish, A., Gerstein, M., Hubner, N., Ingolia, N. T., Kellis, M., Menschaert, G., Moritz, R. L., Ohler, U., Roucou, X., Saghatelian, A., Weissman, J. S., and van Heesch, S. (July, 2022) Standardized Annotation of Translated Open Reading Frames. Nature Biotechnology, 40(7), 994–999.

14. Dunn, J. G. and Weissman, J. S. (2016) Plastid: nucleotide-resolution analysis of next-generation sequencing and genomics data. BMC Genomics, 17(1), 958.

15. Lauria, F., Tebaldi, T., Bernabò, P., Groen, E. J. N., Gillingwater, T. H., and Viero, G. (August, 2018) riboWaltz: Optimization of Ribosome P-site Positioning in Ribosome Profiling Data. PLOS Computational Biology, 14(8), e1006169.

16. Xiao, Z., Huang, R., Xing, X., Chen, Y., Deng, H., and Yang, X. (June, 2018) De Novo Annotation and Characterization of the Translatome with Ribosome Profiling Data. Nucleic Acids Research, 46(10), e61.

17. Clauwaert, J., McVey, Z., Gupta, R., and Menschaert, G. (March, 2023) TIS Transformer: Remapping the Human Proteome Using Deep Learning. NAR genomics and bioinformatics, 5(1), qad021.

18. Dobin, A. and Gingeras, T. R. (2015) Mapping RNA-seq Reads with STAR. Current Protocols in Bioinformatics, 51(1), 11.14.1–11.14.19.

19. Martin, M. (May, 2011) Cutadapt Removes Adapter Sequences from High-Throughput Sequencing Reads. EMB-net.journal, 17(1), 10–12.

20. Choromanski, K., Likhosherstov, V., Dohan, D., Song, X., Gane, A., Sarlos, T., Hawkins, P., Davis, J., Mohiuddin, A., Kaiser, L., Belanger, D., Colwell, L., and Weller, A. (March, 2021) Rethinking Attention with Performers. arXiv:2009.14794 [cs, stat],.

21. Gawron, D., Ndah, E., Gevaert, K., and Van Damme, P. (February, 2016) Positional Proteomics Reveals Differences in N-terminal Proteoform Stability. Molecular Systems Biology, 12(2), 858.

22. Stern-Ginossar, N., Weisburd, B., Michalski, A., Le, V. T. K., Hein, M. Y., Huang, S.-X., Ma, M., Shen, B., Qian, S.-B., Hengel, H., Mann, M., Ingolia, N. T., and Weissman, J. S. (November, 2012) Decoding Human Cytomegalovirus. Science, 338(6110), 1088–1093.

23. Gonzalez, C., Sims, J. S., Hornstein, N., Mela, A., Garcia, F., Lei, L., Gass, D. A., Amendolara, B., Bruce, J. N., Canoll, P., and Sims, P. A. (August, 2014) Ribosome Profiling Reveals a Cell-Type-Specific Translational Landscape in Brain Tumors. Journal of Neuroscience, 34(33), 10924–10936.

24. Rubio, C. A., Weisburd, B., Holderfield, M., Arias, C., Fang, E., DeRisi, J. L., and Fanidi, A. (October, 2014) Transcriptome-Wide Characterization of the eIF4A Signature Highlights Plasticity in Translation Regulation. Genome Biology, 15(10), 476.

25. Werner, A., Iwasaki, S., McGourty, C. A., Medina-Ruiz, S., Teerikorpi, N., Fedrigo, I., Ingolia, N. T., and Rape, M. (September, 2015) Cell-Fate Determination by Ubiquitin-Dependent Regulation of Translation. Nature, 525(7570), 523–527.

26. Tanenbaum, M. E., Stern-Ginossar, N., Weissman, J. S., and Vale, R. D. (August, 2015) Regulation of mRNA Translation during Mitosis. eLife, 4, e07957.

27. Zur, H., Aviner, R., and Tuller, T. (February, 2016) Complementary Post Transcriptional Regulatory Information Is Detected by PUNCH-P and Ribosome Profiling. Scientific Reports, 6(1), 21635.

28. Loayza-Puch, F., Rooijers, K., Buil, L. C. M., Zijlstra, J., F. Oude Vrielink, J., Lopes, R., Ugalde, A. P., van Breugel, P., Hofland, I., Wesseling, J., van Tellingen, O., Bex, A., and Agami, R. (February, 2016) Tumour-Specific Proline Vulnerability Uncovered by Differential Ribosome Codon Reading. Nature, 530(7591), 490–494.

29. Bencun, M., Klinke, O., Hotz-Wagenblatt, A., Klaus, S., Tsai, M.-H., Poirey, R., and Delecluse, H.-J. (April, 2018) Translational Profiling of B Cells Infected with the Epstein-Barr Virus Reveals 51 Leader Ribosome Recruitment through Upstream Open Reading Frames. Nucleic Acids Research, 46(6), 2802–2819.

30. Raj, A., Wang, S. H., Shim, H., Harpak, A., Li, Y. I., Engelmann, B., Stephens, M., Gilad, Y., and Pritchard, J. K. (May, 2016) Thousands of Novel Translated Open Reading Frames in Humans Inferred by Ribosome Footprint Profiling. eLife, 5, e13328.

31. Fang, H., Huang, Y.-F., Radhakrishnan, A., Siepel, A., Lyon, G. J., and Schatz, M. C. (February, 2018) Scikit-Ribo Enables Accurate Estimation and Robust Modeling of Translation Dynamics at Codon Resolution. Cell Systems, 6(2), 180–191.e4.

32. Crappé, J., Ndah, E., Koch, A., Steyaert, S., Gawron, D., De Keulenaer, S., De Meester, E., De Meyer, T., Van Criekinge, W., Van Damme, P., and Menschaert, G. (March, 2015) PROTEOFORMER: Deep Proteome Coverage through Ribosome Profiling and MS Integration. Nucleic Acids Research, 43(5), e29.

33. Zhang, P., He, D., Xu, Y., Hou, J., Pan, B.-F., Wang, Y., Liu, T., Davis, C. M., Ehli, E. A., Tan, L., Zhou, F., Hu, J., Yu, Y., Chen, X., Nguyen, T. M., Rosen, J. M., Hawke, D. H., Ji, Z., and Chen, Y. (November, 2017) Genome-Wide Identification and Differential Analysis of Translational Initiation. Nature Communications, 8(1), 1749.

34. Erhard, F., Halenius, A., Zimmermann, C., L’Hernault, A., Kowalewski, D. J., Weekes, M. P., Stevanovic, S., Zimmer, R., and Dölken, L. (May, 2018) Improved Ribo-seq Enables Identification of Cryptic Translation Events. Nature Methods, 15(5), 363–366.

35. Malone, B., Atanassov, I., Aeschimann, F., Li, X., Großhans, H., and Dieterich, C. (April, 2017) Bayesian Prediction of RNA Translation from Ribosome Profiling. Nucleic Acids Research, 45(6), 2960–2972.

36. Choudhary, S., Li, W., and D. Smith, A. (April, 2020) Accurate Detection of Short and Long Active ORFs Using Ribo-seq Data. Bioinformatics, 36(7), 2053–2059.

