## Supplemental File for "RIBO-former: leveraging ribosome profiling information to improve the detection of translated open reading frames"

### 1. Ribosome profiling processing

A wide selection of data sources is included as part of this study (see Supplementary Table A2). All data is processed following the same steps. All steps have been performed on an server featuring >40 CPU cores, 100GB RAM, and 1 RTX 3090 GPU. This section features multiple code snippets used to run the code.

#### 1.1. Mapping

Cutadapt and STAR are applied for trimming adapters and mapping reads to the genome and transcriptome. Read lengths between 20 and 40 nucleotides are retained. Reads mapping against tRNA/rRNA/sm(o)RNA are filtered out. Ensembl GRCh38-v107 is used as the reference genome and annotation. Supplementary Table A3 lists the total number of reads within the data set at various steps. The data were selected in order to have variation with respect to applied treatments and mapped number of reads.

```
# trim files and perform fastqc
cutadapt -j 20 -m 20 -a $adapter ${dataset}.fastq -o "out/temp/${dataset}_trimmed.fq" > "
    ↪ out/temp/${dataset}_trimmed_report.txt"

# remove rRNA/tRNA/smRNA/smoRNA
STAR --genomeLoad NoSharedMemory --seedSearchStartLmaxOverLread .5 --genomeDir '../../
    ↪ genome/STAR/excl_RNA' --readFilesIn $trimmed --outFilterMultimapNmax 1000 --
    ↪ outFilterMismatchNmax 2 --outFileNamePrefix out/temp/ --runThreadN 20 --
    ↪ outReadsUnmapped Fastx
mv out/temp/Unmapped.out.mate1 $cleaned

# align to genome, output mapping to transcriptome as well
STAR --runThreadN 20 --genomeDir '../../genome/STAR' --genomeLoad NoSharedMemory --
    ↪ readFilesIn $cleaned --outFileNamePrefix out/ --outSAMtype BAM SortedByCoordinate
    ↪ --quantMode TranscriptomeSAM --outSAMattributes MD NH --outFilterMultimapNmax 10
    ↪ --outMultimapperOrder Random --outFilterMismatchNmax 2 --
    ↪ seedSearchStartLmaxOverLread 0.5 --alignEndsType EndToEnd --outWigType bedGraph
```

The data contained various adapters.

```
# benchmarked data sets
SRR1802129 CTGTAGGCACCATCAATAGATCGGAAGAGC
SRR2433794 TGGAATTCTCGG
SRR2732970 CTGTAGGCACCATCAATAGATCGGAAGAGCACAC
SRR2733100 CTGTAGGCACCATCAATAGATCGGAAGAGCACAC
SRR2954800 TGGAATTCTCGG
SRR8449577 AGATCGGAAGAGC
SRR9113067 AGATCGGAAGAGC
SRR11005875 AGATCGGAAGAGC

# biological replicates
SRR9113068 AGATCGGAAGAGC
SRR9113069 AGATCGGAAGAGC
SRR8449578 AGATCGGAAGAGC

# pre-training machine learning model
SRR592960 CTGTAGGCACCATCAATTCG
SRR1562539 CTGTAGGCACCATCAATCGACGTATCTCGTATGCCGTCTTCT
SRR1573939 CTGTAGGCACCATCAATTCGTATGCCGTC
SRR1610244 CTGTAGGCACCATTAATAGATC
SRR1976443 CTGTAGGCACCATCAATTCGTA
SRR2536856 CTGTAGGCACCATCAATTCGTATGCCGTC
SRR2873532 TGGAATTCTCGGGTGCCAAGGAACTCCAGTCACCG
```

```
SRR3575904 AGATCGGAAGAGC
```

### 2. RIBO-former

#### 2.1. Data processing

Data loading for RIBO-former is achieved by storing data in the hierarchical data format version 5 (*hdf5*). Using Python, the ribosome reads mapped to the transcriptome are stored by transcript. The generated *bam* files are parsed using Python and data is stored to the *hdf5* format. Data is aggregated by the total number of reads aligned by their 5' position for every read length and transcript position. Transcript matrices are loaded from the *hdf5* files by a *pytorch* data loader object and used as inputs to the model.

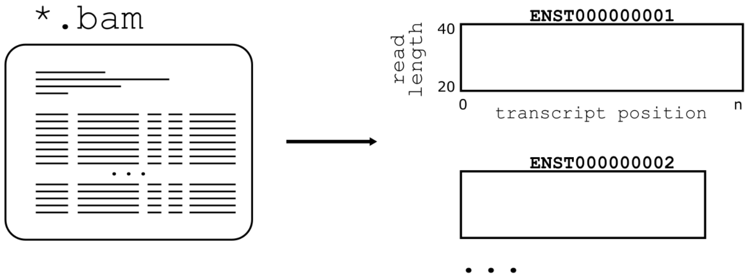


#### 2.2. Input embedding strategies

The transformer architecture takes full transcript regions as input and provides a prediction along each position of the input range. No sequence information is processed. No ORFs are identified as a pre-processing step. Transformer networks use vector representations of mapped reads at each nucleotide position as input tokens. As part of this research, different approaches were explored to create input vector representations from the mapped ribosome profiling data. For all instances, read counts are normalized for each transcript. This ensures the numerical stability of the inputs. Supplementary Figures A8 illustrates both strategies evaluated as part of this paper. Supplementary Section 2.3 displays the number of model parameters in each module of the network. See the main manuscript on the implementation of each strategy.

#### 2.3. Model architecture

The architectural framework is identical to that of TIS transformer Clauwaert et al. [2023], a transformer model used for predicting translation initiation sites using transcript sequence information. The transformer structure features multiple layers with multiple attention heads per layer. These are identical in structure but feature unique trainable model parameters. The outputs of the transformer module are sent to a set of fully connected layers to obtain a binary output at each input position. Notwithstanding the size of the data set and overall high computational requirements of transformer architectures, model optimization **from scratch** is possible on a single RTX 3090 and converges after ca. 6 hours due to the relative shallowness of the transformer architecture as compared to many language-learning transformers.

**Algorithm 1** RIBO-former network architecture. Given are the different layers, their respective dimensions as defined by their hyperparameter names, the dimensions for RIBO-former (Table A5), and the resulting total weights. The bias term applied in each node is included and marked with italics.

**RIBO-former** | **211,964**
- **Ribosome Read Count** | **21,546**
  - **Linear** | $1 \times \text{dim}$ | $1 \times 42 + \mathit{42}$ | **84**
  - **Linear** | $\text{dim} \times \text{dim} * 6$ | $42 \times 252 + \mathit{252}$ | **10,836**
  - **Linear** | $\text{dim} * 6 \times \text{dim}$ | $252 \times 42 + \mathit{42}$ | **10,626**
- **Ribosome Read Count Embedding** | $1 \times \text{dim}$ | $1 \times 42$ | **42**
- **Ribosome Read Length Embedding** | read lengths $\times$ dim | $21 \times 42$ | **882**
- **Positional Embedding** | fixed positional embeddings | **0**
- **Performer** | **185,712**
  - **Layer** ($\times$ depth | 6) | **30,952**
    - **Layer norm** | $\text{dim} \times 2$ | $42 \times 2 + \mathit{2}$ | **86**
    - **Attention head** ($\times$ n_head | 6) | **2,064**
      - $\mathbf{W_Q}$ | dim $\times$ dim_head | $42 \times 16 + \mathit{16}$ | **688**
      - $\mathbf{W_K}$ | dim $\times$ dim_head | $42 \times 16 + \mathit{16}$ | **688**
      - $\mathbf{W_V}$ | dim $\times$ dim_head | $42 \times 16 + \mathit{16}$ | **688**
    - $\mathbf{W_o}$ | dim_head $*$ n_head $\times$ dim | $96 \times 42 + \mathit{42}$ | **4,074**
    - **Layer norm** | $\text{dim} \times 2$ | $42 \times 2 + \mathit{2}$ | **86**
    - **Linear** | $\text{dim} \times \text{dim} * 4$ | $42 \times 168 + \mathit{168}$ | **7,224**
    - **Linear** | $\text{dim} * 4 \times \text{dim}$ | $168 \times 42 + \mathit{42}$ | **7,098**
- **Linear** | $\text{dim} \times \text{dim} * 2$ | $42 \times 84 + \mathit{84}$ | **3,612**
- **Linear** | $\text{dim} \times 2$ | $84 \times 2 + \mathit{2}$ | **170**

#### 2.3.1. Attention

Custom attention strategies can be performed by the attention heads independent of the number of weights utilized to calculate the **Q**, **K**, **V** matrices. In this model, full attention is calculated through the Fast Attention Via Positive Orthogonal Random Features (FAVOR+) algorithm (Figure A1: left) Choromanski et al. [2021]. These allow full attention, where all inputs along the transcript are included by the attention head. In contrast, local attention restricts the attention matrix to only neighboring positions. Local attention is implemented by dividing the attention matrix in smaller blocks on which full attention is calculated (Figure A1: right). Three blocks around the evaluated input are calculated. These local attention heads do not apply the FAVOR+ algorithm and use rotary positional embeddings Su et al. [2022]. The block size of the local attention heads is referred to under the 'attention scheme' columns of Supplementary Table A5.

### 3. Training and Evaluation

This study explores the use of transformer models to detect translated coding sequences using ribosome profiling data. This is achieved by detecting translated initiation sites, constituting a binary-classification task. Model evaluations follow a standard deep learning set-up featuring a training, validation and test set. Data is grouped according to chromosomes to prevent identical profiles of ribosome reads, possible due to the existence of transcript isoforms, being separated between the training, validation or test set. The data used for the hyperparameter and input strategy selection are chromosomes 3, 4, 5, 6, 8, 9, 10, 11, 12, 13, 15, 16, 17, 18, 20, 21, 22, X, and Y for the training set, chromsomes 2 and 14 for the validation set, and chromosomes 1, 7, 13, and 19 for the test set. The data used for the pre-training strategy selection and benchmarking with previous tools feature two sets (folds) in order to cover the full transcriptome (within the test set). The first fold has chromosomes 3, 5, 7, 11, 13, 15, 19, 21, and X for the training set, chromosomes 1, 9, and 17 for the validation set, and chromosomes 2, 4, 6, 8, 10, 12, 14, 16, 18, 20, 22, and Y for the test set. The second fold has chromosomes 2, 6, 8, 10, 14, 16, 18, 22, and Y for the training set, chromosomes 4, 12, and 20 for the validation set, and chromosomes 1, 3, 5, 7, 9, 11, 13, 15, 17, 19, 21, and X for the test set. The cross-entropy is used to determine the loss. For all set-ups described in this paper, a learning rate of $10^{-3}$ is applied. The loss on the validation set indicates the optimal point of the model fit and is used for model selection (i.e. early stopping) to prevent overfitting on the training set. All reported performances are obtained on the test set.

#### 3.1. Positive set and performance

To train and evaluate the model, translation initiation sites have been labeled using the Ensembl GRCh38v107 annotation. The full transcriptome constitutes 251,121 transcript regions, with a total of 431,011,438 positions. The aim of the paper is to predict the translatome, i.e. the mRNA that is being translated. As explained in the main manuscript, we have decided to use all known translation initiation sites as the positive set. We do this as filtering the positive set has no advantage but does incur an unnecessary factor of noise (selection of thresholds, and thus inclusion of existing translation initiation sites into the positive set is pseudo-arbitrary). Sub-setting the input data or altering the labels can furthermore hinder utility and benchmarking efforts. While including all known TISs to the positive set does effect the maximum performance a tool can achieve using a certain data set, as an unknown amount of coding sequences are not being translated or have simply not been captured by the ribosome profiling experiment, it does not hinder comparison between different tools. A higher performance score indicates one approach to have reconstructed a larger part of the proteome–and thus translatome–using ribosome profiling data, supporting its superiority over the other.

#### 3.2. Hyperparameter selection

Hyperparameter optimization is performed to identify the optimal model architecture for this learning problem. The data set featuring the highest number of mapped reads (SRR2733100) is selected to perform the hyperparameter selection on, as this ensures the complexity of the task is properly represented by the data (i.e. less reads feature a less complex task). No individual hyperparameters were observed to be more effective than others in improving performances. However, a correlation exists between the total number of model parameters and model performance. Eight unique architectures have been evaluated featuring varying configurations for the size of the hidden dimension, number of layers, number of attention heads per layer and the dimension of the attention head itself (Supplementary Table A5). Supplementary Figure A10 shows the validation loss at different epochs of the various architectures.

#### 3.3. Input token strategy

The input token strategies follows the same data allocations as the hyperparameter selection. To evaluate strategy A, reads were mapped by their 5' reads without offsets, and with offsets calculated by Plastid and RiboWaltz. Plastid and RiboWaltz have been selected for A/P-site offset calling as they are both popular methods specifically created for this task. Package versions are those listed as 'Last update' in Table A1. Scripts are executed following our custom folder structure.

**Plastid**

```
reformat_transcripts --annotation_files genome/Homo_sapiens.GRCh38.107.gff3 --
    ↪ annotation_format GFF3 --output_format GTF2 genome/Homo_sapiens.GRCh38.107.gtf2
metagene generate genome/plastid/ --landmark cds_start --annotation_files genome/
    ↪ Homo_sapiens.GRCh38.107.gtf2
psite genome/plastid/_rois.txt ribo/${dataset}/out/plastid/ --min_length 20 --max_length
    ↪ 41 --require_upstream --count_files ribo/${dataset}/out/genome/${dataset}_aligned.
    ↪ bam
```

**RiboWaltz**

```
library(riboWaltz)

metadata <- read.table('ribo/metadata.txt', header = FALSE, sep = "", dec = ".")
annotation_db <- create_annotation('genome/Homo_sapiens.GRCh38.107.gtf')
for (i in metadata$V1){
   reads_list <- bamtolist(bamfolder=sprintf("ribo/%s/out/", i), annotation=annotation_db
       ↪ )
   filtered_list <- length_filter(data=reads_list, length_filter_mode="custom", length_
       ↪ range=20:40)
   psite_offset <- psite(filtered_list)
   dir.create(sprintf("ribo/%s/out/ribowaltz", i))
   write.table(psite_offset, sprintf("ribo/%s/out/ribowaltz/riboWaltz_offsets.csv", i),
       ↪ sep="\t")
}
```

#### 3.4. (Pre-)training

Training a model that is already optimized on ribosome data shows two important advantages as compared to training a new model from scratch: faster convergence of the validation loss, substantially reducing optimization times, and improved performances. This indicates that some correlations are shared between ribosome data sets.

Model pre-training follows the same data groupings to ensure models are exposed to the same transcripts during training at any stage. This is an additional measure as it is possible for the model to identify transcripts by their lengths and directly learn the location of the transcription initiation in a case of model overfitting. Two settings of model pre-training have been evaluated using eight additional data sets (Table A2). Given the results of the previous section, strategy B is the default method for computing input tokens.

To apply RIBO-former for mapping translation initiation sites on the transcriptome, it is necessary to train multiple models from which different parts of the transcriptome are excluded during training or model selection (i.e. training and validation set). Selecting a larger fraction of the transcriptome for training can result in better generalization due to the larger amount of data the model is fit on. However, the size of the training and validation set affects that of the test set, where more models need to be optimized as the fraction of the test set decreases in order to cover the full transcriptome. In this study, we have chosen to evaluate RIBO-former using the minimum training and validation set available, splitting the data in half, requiring only two models to cover the full transcriptome. We expect the tool to be used as such as it constitutes the least amount of computing time, and have therefore evaluated RIBO-former using the minimum of two folds. Identical data allocations are used for pre-training and training of the models.

#### 3.5. Benchmark

Several tool exist that utilize ribosome profiling data in various ways to delineate translated open reading frames (Table A1. Specifically, we discuss the output and results from RIBO-former with those of PRICE Erhard et al. [2018], Rp-Bp Malone et al. [2017], RiboTISH Zhang et al. [2017], and RiboCode Xiao et al. [2018].

Several differences exist between the approach of RIBO-former and discussed methods. RIBO-former processes ribosome mapping profiles for each transcript to predict the presence of translation initiation sites at each position. No information of the sequence or valid open reading frames ensures that the model is protected from learning certain biases that are often understood to be present today's genome annotation, including the under-representation of non-canonical start codons and small ORFs. However, the exclusion of these types of information constitutes a harder learning problem. While it is possible to only evaluate the model predictions for specific start codons, for example, to get an idea of the model performance for those sites, this still constitutes a disadvantage as compared to an approach that has sequence information as an input feature.

Post-processing steps performed by existing tools are omitted as these obscure the predictive capability of the algorithm. For example, multiple methods allow you to filter down the results to only include the longest possible ORF on a transcript featuring an ATG start codon.

**PRICE**

```
gedi -e IndexGenome -s genome/Homo_sapiens.GRCh38.dna.primary_assembly.fa -a genome/
    ↪ Homo_sapiens.GRCh38.107.gtf -f genome/price -nobowtie -nostar -nokallisto
gedi -e Price -reads ribo/${dataset}/out/genome/${dataset}_aligned.bam -genomic
    ↪ Homo_sapiens.GRCh38.107 -prefix ribo/${dataset}/out/price/ -progress -plot
```

**Rp-Bp**

```
prepare-rpbp-genome ../scripts/benchmark/rpbp_full.yml --star-options "--
    ↪ genomeSAindexNbases 10" --mem 10G --num-cpus 4 --logging-level INFO --log-file
    ↪ genome/rpbp/rpbp-genome.log
run-all-rpbp-instances ribo/${dataset}/out/rpbp/rpbp.yml --num-cpus 30 --logging-level
    ↪ INFO --mem 50G
```

**RiboCode**

```
prepare_transcripts -g genome/Homo_sapiens.GRCh38.107.gtf -f genome/Homo_sapiens.GRCh38.dna.primary_assembly.fa -o genome/ribocode
metaplots -a genome/ribocode -m 20 -M 40 -r ribo/${dataset}/out/${dataset}_aligned_tran.bam -o ribo/${dataset}/out/ribocode/
RiboCode -a genome/ribocode -c ribo/${dataset}/out/ribocode/_pre_config.txt -l no -g -o ribo/${dataset}/out/ribocode/
```

#### RiboTish

```
ribotish quality -b ribo/${dataset}/out/genome/${dataset}_aligned.bam -g genome/Homo_sapiens.GRCh38.107.gtf -f ribo/${dataset}/out/ribotish/quality.pdf -r ribo/${dataset}/out/ribotish/offset.txt -o ribo/${dataset}/out/ribotish/quality.txt -l 20,41
ribotish predict -b ribo/${dataset}/out/genome/${dataset}_aligned.bam -g genome/Homo_sapiens.GRCh38.107.gtf -f genome/Homo_sapiens.GRCh38.dna.primary_assembly.fa -o ribo/${dataset}/out/ribotish/orfs.txt --ribopara ribo/${dataset}/out/ribotish/offset.txt
```

#### Ribotricer

```
ribotricer prepare-orfs --gtf genome/Homo_sapiens.GRCh38.107.gtf --fasta genome/Homo_sapiens.GRCh38.dna.primary_assembly.fa --prefix genome/ribotricer/ribo
ribotricer detect-orfs --bam ribo/${dataset}/out/genome/${dataset}_aligned.bam --ribotricer_index genome/ribotricer/ribo_candidate_orfs.tsv --prefix ribo/${dataset}/out/ribotricer/
```

### 4. Supplementary Tables

Table A1: **Various tools designed to detect expressed coding sequences using ribosome profiling data.** Methods with * require RNA-seq data. The list is non-exhaustive.

| | | Calling | | | | |
|---|---|---|---|---|---|---|
| **Method** | **Author** | **A/P-site** | **ORF** | **Year** | **Last update** | **Language** |
| Ribotricer | Choudhary et al. [2020] | Yes | Yes | 2019 | 1.3.3 (2023) | Python |
| Ribodeblur | Ahmed et al. [2019] | Yes | No | 2018 | 2018 | Python |
| RiboWaltz | Lauria et al. [2018] | Yes | No | 2018 | 1.2.0 (2021) | R |
| RibORF | Ji [2018] | Yes | Yes | 2018 | 2.0 (2022) | Perl |
| RiboCode | Xiao et al. [2018] | Yes | Yes | 2018 | 1.2.15 (2022) | Python |
| RiboWave | Xu et al. [2018] | Yes | Yes | 2018 | 2018 | Python |
| Scikit-ribo* | Fang et al. [2018] | No | Yes | 2018 | 2018 | Python |
| Rp-Bp | Malone et al. [2017] | Yes | Yes | 2017 | 3.0.1 (2023) | Python |
| RiboTISH | Zhang et al. [2017] | Yes | Yes | 2017 | 0.2.7 (2021) | Python |
| Plastid | Dunn and Weissman [2016] | Yes | No | 2016 | 0.6.1 (2022) | Python |
| PRICE (Gedi) | Erhard et al. [2018] | No | Yes | 2016 | 1.0.5 (2022) | Python |
| SPECtre | Chun et al. [2016] | No | Yes | 2016 | 1.0.0 (2018) | R/Python |
| riboHMM* | Raj et al. [2016a] | No | Yes | 2016 | 2016 | Python |
| RiboProfiling | Popa et al. [2016] | Yes | Yes | 2016 | 1.28.0 (2022) | R |
| RiboTaper* | Calviello et al. [2016] | No | Yes | 2015 | 1.3 (2016) | R |
| ORF-RATER | Fields et al. [2015] | No | Yes | 2015 | 2018 | Python2.7 |
| PROTEOFORMER | Crappé et al. [2015] | Plastid | Yes | 2015 | 2.0 (2022) | Python2.7/Perl |

Table A2: **Data sets and their corresponding publications used in this study.** Generally, only a single data set (SRR entry) is used from each study. ER: estrogen receptor, iPSCs: induced pluripotent stem cells, hESC: embrionic stem cells, HCMV: human cytomegalovirus, RPE: retinal pigment epithelium. EBV: epstein-bar virus

| **Author** | **Year** | **SRR entry** | **Treatment** | **Tissue** |
|---|---|---|---|---|
| **Benchmarking** | | | | |
| Ji et al. [2015] | 2015 | SRR1802129 | Cycloheximide | MCF10A-ER-Src |
| Calviello et al. [2016] | 2016 | SRR2433794 | Cycloheximide | HEK293 |
| Gawron et al. [2016] | 2016 | SRR2732970 | Lactimidomycin | Jurkat |
| Gawron et al. [2016] | 2016 | SRR2733100 | Cycloheximide | Jurkat |
| Raj et al. [2016b] | 2016 | SRR2954800 | Harringtonine | Lymphoblastoid |
| Martinez et al. [2020] | 2020 | SRR8449577 | Cycloheximide | HeLa S3 |
| Chen et al. [2020] | 2020 | SRR9113067 | – | iPSC |
| Gaertner et al. [2020] | 2020 | SRR11005875 | Cycloheximide | Pancreatic hESC |
| **Pre-training** | | | | |
| Stern-Ginossar et al. [2012] | 2012 | SRR592960 | Lactimidomycin | HCMV foreskin fibroblasts |
| Gonzalez et al. [2014] | 2014 | SRR1562539 | Cycloheximide | Brain |
| Rubio et al. [2014] | 2014 | SRR1573939 | Cycloheximide | MDA-MB-231 |
| Werner et al. [2015] | 2015 | SRR1610244 | Cycloheximide | hES |
| Tanenbaum et al. [2015] | 2015 | SRR1976443 | Cycloheximide | RPE |
| Zur et al. [2016] | 2016 | SRR2536856 | Cycloheximide | HeLa S3 |
| Loayza-Puch et al. [2016] | 2016 | SRR2873532 | Chloramphenicol | PC-3 |
| Bencun et al. [2018] | 2018 | SRR3575904 | Harringtonine | CD19 EBV |

Table A3: **Read counts at different stages of the mapping process**. The adapters are trimmed from the reads (+trim) after which reads mapping to sn(o)RNA/tRNA/rRNA (xRNA) are filtered out (+trim-xRNA). The duplication rates are obtained as a function of reads mapped to the genome.

| | | | | | Mapped | |
|---|---|---|---|---|---|---|
| **Data set** | **Raw** | **+ trim** | **+trim −xRNA** | **xRNA** | **Genome** | **Dupl. rate** |
| **Benchmarking** | | | | | | |
| SRR1802129 | 2.06E+07 | 2.05E+07 | 2.22E+06 | 89.14% | 2.84E+06 | 1.69 |
| SRR2433794 | 3.20E+07 | 3.19E+07 | 2.75E+07 | 13.81% | 3.56E+07 | 1.50 |
| SRR2732970 | 3.78E+08 | 3.05E+08 | 1.42E+08 | 53.29% | 1.95E+08 | 3.03 |
| SRR2733100 | 3.89E+08 | 3.63E+08 | 1.88E+08 | 48.28% | 2.46E+08 | 4.33 |
| SRR2954800 | 6.61E+06 | 6.61E+06 | 4.03E+06 | 39.12% | 3.87E+06 | 2.23 |
| SRR8449577 | 6.50E+07 | 6.42E+07 | 1.07E+07 | 83.33% | 1.19E+07 | 1.78 |
| SRR9113067 | 1.50E+07 | 1.25E+07 | 5.34E+06 | 57.20% | 6.04E+06 | 2.01 |
| SRR11005875 | 3.01E+07 | 3.00E+07 | 9.87E+06 | 67.12% | 9.75E+06 | 1.39 |
| **Pre-training** | | | | | | |
| SRR592960 | 4.62E+07 | 4.48E+07 | 9.14E+06 | 79.59% | 4.69E+06 | 1.30 |
| SRR1562539 | 5.65E+07 | 5.18E+07 | 1.98E+07 | 61.88% | 1.43E+07 | 9.78 |
| SRR1573939 | 3.88E+08 | 3.70E+08 | 5.95E+07 | 83.92% | 8.12E+07 | 1.34 |
| SRR1610244 | 4.78E+07 | 4.14E+07 | 1.22E+07 | 70.48% | 1.43E+07 | 1.56 |
| SRR1976443 | 1.92E+08 | 1.84E+08 | 3.35E+07 | 81.78% | 4.40E+07 | 1.89 |
| SRR2536856 | 6.65E+07 | 5.17E+07 | 1.29E+07 | 75.07% | 1.74E+07 | 1.82 |
| SRR2873532 | 3.37E+07 | 3.35E+07 | 1.53E+07 | 54.28% | 1.93E+07 | 1.65 |
| SRR3575904 | 5.61E+07 | 5.61E+07 | 6.35E+06 | 88.69% | 8.47E+06 | 1.83 |
| **Replicates** | | | | | | |
| SRR9113068 | 2.43E+07 | 2.14E+07 | 9.44E+06 | 55.86% | 1.18E+07 | 2.48 |
| SRR9113069 | 2.24E+07 | 1.96E+07 | 1.08E+07 | 44.97% | 1.22E+07 | 2.35 |
| SRR8449578 | 1.15E+08 | 1.12E+08 | 2.13E+07 | 81.08% | 2.47E+07 | 1.64 |

Table A4: **A/P-offsets determined by various tools evaluated during this study.** The listed tools are Plastid (P), RiboWaltz (W), RiboCode (C), and RiboTISH (T)

| | **SRR1802129** | | | | | **SRR2433794** | | | | | **SRR2732970** | | | | | **SRR2733100** | | | | |
|---|---|---|---|---|---|---|---|---|---|---|---|---|---|---|---|---|---|---|---|---|
| | **%** | **P** | **W** | **C** | **T** | **%** | **P** | **W** | **C** | **T** | **%** | **P** | **W** | **C** | **T** | **%** | **P** | **W** | **C** | **T** |
| **20** | 0.2 | 13 | 7 | - | - | 0.1 | 13 | 13 | - | - | 1.3 | 2 | 12 | - | - | 0.8 | 2 | 12 | - | - |
| **21** | 0.2 | 13 | 9 | 5 | 17 | 0.2 | 13 | 13 | - | - | 1.6 | 3 | 12 | - | 12 | 1.0 | 3 | 12 | 12 | 12 |
| **22** | 0.2 | 13 | 9 | - | - | 0.4 | 5 | 12 | - | - | 1.7 | 12 | 12 | - | - | 1.0 | 3 | 12 | - | - |
| **23** | 0.2 | 13 | 7 | - | 7 | 0.7 | 6 | 12 | - | 12 | 2.2 | 12 | 12 | - | - | 1.2 | 5 | 12 | - | - |
| **24** | 0.3 | 13 | 8 | - | 8 | 1.0 | 7 | 12 | - | - | 2.2 | 12 | 12 | - | 12 | 1.4 | 5 | 12 | - | 12 |
| **25** | 0.4 | 13 | 9 | - | 12 | 2.4 | 9 | 12 | 12 | 12 | 2.3 | 12 | 12 | - | - | 1.5 | 12 | 12 | - | - |
| **26** | 1.7 | 50 | 10 | - | - | 5.2 | 9 | 12 | - | 12 | 2.5 | 12 | 12 | - | - | 1.8 | 12 | 12 | - | - |
| **27** | 9.6 | 12 | 12 | - | 11 | 8.7 | 10 | 12 | - | 12 | 3.1 | 12 | 12 | 12 | 12 | 2.5 | 12 | 12 | 12 | 12 |
| **28** | 36.3 | 12 | 12 | 12 | 12 | 19.7 | 12 | 12 | - | 12 | 4.8 | 12 | 12 | 12 | 12 | 4.3 | 12 | 12 | - | 12 |
| **29** | 32.0 | 12 | 13 | - | 12 | 37.4 | 12 | 12 | 12 | 12 | 14.5 | 12 | 12 | 12 | 12 | 13.3 | 12 | 12 | 12 | 12 |
| **30** | 9.0 | 23 | 15 | - | 13 | 21.8 | 12 | 12 | 12 | 12 | 30.0 | 12 | 12 | 12 | 12 | 27.7 | 12 | 12 | 12 | 12 |
| **31** | 1.7 | 13 | 14 | - | - | 2.1 | 13 | 13 | - | - | 21.6 | 12 | 12 | 12 | 12 | 25.0 | 12 | 12 | - | 12 |
| **32** | 1.4 | 13 | 16 | - | - | 0.3 | 50 | 11 | - | - | 8.3 | 12 | 13 | 12 | 12 | 12.5 | 13 | 13 | - | - |
| **33** | 2.5 | 13 | 17 | - | - | 0.1 | 13 | 12 | - | 15 | 2.5 | 13 | 13 | - | - | 4.0 | 5 | 13 | - | - |
| **34** | 2.5 | 13 | 19 | - | - | 0.0 | 13 | 14 | - | - | 0.8 | 12 | 13 | - | - | 1.0 | 5 | 13 | - | - |
| **35** | 1.4 | 13 | 18 | - | - | 0.0 | 13 | 13 | - | - | 0.3 | 13 | 13 | - | - | 0.4 | 13 | 14 | - | - |
| **36** | 0.1 | 13 | 20 | - | - | 0.0 | 13 | 15 | - | - | 0.2 | 12 | 12 | - | - | 0.2 | 13 | 12 | - | - |
| **37** | 0.1 | 13 | 21 | - | - | 0.0 | 13 | 12 | - | - | 0.1 | 50 | 12 | - | - | 0.2 | 13 | 14 | - | - |
| **38** | 0.1 | 13 | 22 | - | - | 0.0 | 13 | 12 | - | - | 0.1 | 13 | 12 | - | - | 0.1 | 13 | 12 | - | - |
| **39** | 0.1 | 13 | 23 | - | - | 0.0 | 13 | 12 | - | - | 0.0 | 50 | 12 | - | - | 0.1 | 13 | 14 | - | - |
| **40** | 0.0 | 13 | 24 | - | - | 0.0 | 13 | 12 | - | - | 0.0 | 13 | 12 | - | - | 0.0 | 13 | 13 | - | - |

| | **SRR2954800** | | | | | **SRR8449577** | | | | | **SRR9113067** | | | | | **SRR11005875** | | | | |
|---|---|---|---|---|---|---|---|---|---|---|---|---|---|---|---|---|---|---|---|---|
| | **%** | **P** | **W** | **C** | **T** | **%** | **P** | **W** | **C** | **T** | **%** | **P** | **W** | **C** | **T** | **%** | **P** | **W** | **C** | **T** |
| **20** | 0.0 | 13 | 7 | - | 9 | 1.0 | 5 | 12 | - | - | 4.6 | 3 | 12 | - | 12 | 0.1 | 13 | 12 | - | 12 |
| **21** | 0.0 | 13 | 11 | - | 8 | 1.6 | 5 | 12 | - | 12 | 6.8 | 12 | 12 | 12 | 12 | 0.4 | 13 | 12 | - | - |
| **22** | 0.1 | 11 | 11 | - | - | 1.6 | 6 | 12 | 12 | 12 | 4.7 | 29 | 13 | - | - | 0.8 | 8 | 12 | - | - |
| **23** | 0.1 | 13 | 12 | - | - | 1.7 | 7 | 12 | - | - | 3.2 | 3 | 12 | - | 12 | 1.3 | 6 | 12 | - | 12 |
| **24** | 0.6 | 5 | 11 | - | 11 | 2.0 | 8 | 12 | - | - | 3.1 | 3 | 12 | - | 12 | 1.7 | 7 | 12 | - | - |
| **25** | 1.4 | 6 | 12 | - | - | 2.9 | 9 | 12 | 12 | 12 | 3.5 | 12 | 12 | - | 12 | 2.1 | 8 | 12 | - | - |
| **26** | 2.1 | 7 | 11 | - | - | 6.3 | 11 | 12 | - | - | 4.3 | 3 | 12 | - | 12 | 3.1 | 9 | 12 | - | 12 |
| **27** | 3.3 | 11 | 11 | 8 | 11 | 15.7 | 12 | 12 | - | - | 5.4 | 12 | 12 | - | 12 | 5.3 | 11 | 12 | - | - |
| **28** | 4.7 | 11 | 11 | - | - | 43.2 | 12 | 12 | 12 | 12 | 8.8 | 12 | 12 | - | 12 | 23.5 | 12 | 12 | 12 | 12 |
| **29** | 10.5 | 11 | 11 | 11 | 11 | 19.2 | 12 | 12 | 12 | 12 | 17.4 | 12 | 12 | 12 | 12 | 45.7 | 12 | 12 | 12 | 12 |
| **30** | 26.9 | 11 | 12 | - | 12 | 2.5 | 6 | 12 | 12 | 12 | 19.0 | 12 | 12 | - | 12 | 13.7 | 12 | 12 | 12 | 12 |
| **31** | 21.0 | 12 | 12 | - | 12 | 0.9 | 13 | 12 | 12 | 12 | 11.5 | 13 | 13 | - | 12 | 1.5 | 50 | 12 | - | - |
| **32** | 9.8 | 12 | 12 | - | 12 | 0.5 | 13 | 12 | - | 12 | 4.9 | 10 | 13 | - | - | 0.3 | 13 | 12 | - | - |
| **33** | 5.9 | 11 | 12 | - | - | 0.4 | 13 | 12 | - | - | 2.0 | 13 | 11 | - | - | 0.2 | 13 | 12 | - | - |
| **34** | 4.5 | 12 | 12 | - | - | 0.2 | 13 | 13 | 12 | - | 0.5 | 13 | 14 | - | - | 0.1 | 13 | 12 | - | - |
| **35** | 3.4 | 13 | 12 | - | - | 0.1 | 13 | 12 | - | - | 0.2 | 13 | 12 | - | - | 0.0 | 13 | 12 | - | 12 |
| **36** | 2.0 | 50 | 12 | - | - | 0.1 | 13 | 14 | - | - | 0.1 | 13 | 11 | - | - | 0.0 | 13 | 10 | - | - |
| **37** | 1.2 | 13 | 11 | - | - | 0.0 | 13 | 11 | - | - | 0.1 | 13 | 11 | - | - | 0.0 | 13 | 18 | - | - |
| **38** | 1.0 | 13 | 13 | - | - | 0.0 | 13 | 12 | - | - | 0.1 | 13 | 12 | - | - | 0.0 | 13 | 12 | - | - |
| **39** | 1.1 | 13 | 11 | - | - | 0.0 | 13 | 12 | - | - | 0.0 | 13 | 12 | - | - | 0.0 | 13 | 12 | - | - |
| **40** | 0.3 | 13 | 13 | - | - | 0.0 | 13 | 12 | - | - | - | 13 | 12 | - | - | 0.0 | 13 | 12 | - | - |

Table A5: **Eight model architectures used for hyperparameter tuning selection.** For each set-up, a model is trained to detect translation initiation sites using ribosome profiling data. Hyperparameter selection is based on the minimum loss on the validation set. Hyperparameter tuning is performed on SRR2733100, featuring the highest read depth of all evaluated data sets. Supplementary Figure A10 displays the validation loss curves for each of the listed architectures.

| | **Hidden** | | **Attention head** | | **Attention scheme** | | | |
|---|---|---|---|---|---|---|---|---|
| **ID** | **state dim.** | **Depth** | **Heads** | **Head dim.** | **Local** | **Full** | **Model parameters** | **Val. loss** ($\times 10^{-3}$) |
| 1 | 24 | 5 | 6 | 12 | 4 | 2 | 81K | 1.099 |
| 2 | 30 | 6 | 6 | 16 | 4 | 2 | 129K | 1.105 |
| 3 | 30 | 8 | 6 | 24 | 4 | 2 | 215K | 1.108 |
| **4** | **42** | **6** | **6** | **16** | **4** | **2** | **211K** | **1.095** |
| 5 | 42 | 8 | 6 | 24 | 4 | 2 | 339K | 1.099 |
| 6 | 48 | 6 | 8 | 16 | 5 | 3 | 297K | 1.096 |
| 7 | 48 | 8 | 8 | 24 | 5 | 3 | 484K | 1.110 |
| 8 | 50 | 10 | 10 | 24 | 6 | 4 | 525K | 1.102 |

Table A6: **RIBO-former performances for different input token strategies and data sets**. Scores are calculated on the test set after selection of the model with the minimum validation loss (See Supplementary Figure A11). For each data set and strategy, the cross-entropy loss ($\times 10^3$), area under the receiver operating characteristic curve (ROC), and area under the precision-recall curve (PR) are given. Results indicate the relevance of read length information for the prediction of translation initiation sites using ribosome profiling data, especially for data sets featuring a higher read depth (see Supplementary Table A3). All strategies are evaluated using the same model architecture and training/validation data (Architecture 4, see Supplementary Table A5, Supplementary Figure A10), Strategy A generates input tokens utilizing read count information for every position of the transcript. Strategy A includes mappings generated by taking the 5' position of every read, and offsetting reads based on read length utilizing two different tools (Plastid, RiboWaltz). Strategy B includes information on both the positions and read lengths of the mapped reads.

| | | **SRR1802129** | | | **SRR2433794** | | | **SRR2732970** | | | **SRR2733100** | | |
|---|---|---|---|---|---|---|---|---|---|---|---|---|---|
| | **Position** | **Loss** | **ROC** | **PR** | **Loss** | **ROC** | **PR** | **Loss** | **ROC** | **PR** | **Loss** | **ROC** | **PR** |
| **A** | 5' | 1.71 | 0.938 | 0.0211 | 1.48 | 0.965 | 0.0937 | 1.40 | 0.963 | 0.161 | 1.38 | 0.965 | 0.161 |
| **A** | Plastid | 1.74 | 0.935 | 0.0144 | 1.49 | 0.964 | 0.0825 | 1.39 | 0.965 | 0.156 | 1.41 | 0.964 | 0.145 |
| **A** | RiboWaltz | 1.7 | 0.941 | 0.0205 | 1.47 | 0.966 | 0.0908 | 1.4 | 0.964 | 0.154 | 1.4 | 0.964 | 0.151 |
| **B** | 5' | **1.69** | 0.945 | 0.0217 | **1.44** | 0.968 | 0.104 | **1.31** | 0.969 | 0.211 | **1.3** | 0.97 | 0.217 |

| | | **SRR2954800** | | | **SRR8449577** | | | **SRR9113067** | | | **SRR11005875** | | |
|---|---|---|---|---|---|---|---|---|---|---|---|---|---|
| | **Position** | **Loss** | **ROC** | **PR** | **Loss** | **ROC** | **PR** | **Loss** | **ROC** | **PR** | **Loss** | **ROC** | **PR** |
| **A** | 5' | 1.69 | 0.935 | 0.0394 | 1.55 | 0.955 | 0.0721 | 1.7 | 0.943 | 0.0178 | 1.48 | 0.967 | 0.0727 |
| **A** | Plastid | 1.72 | 0.931 | 0.0309 | 1.57 | 0.954 | 0.0579 | 1.7 | 0.942 | 0.0161 | 1.5 | 0.966 | 0.0721 |
| **A** | RiboWaltz | 1.71 | 0.932 | 0.0324 | 1.56 | 0.955 | 0.064 | 1.7 | 0.943 | 0.0171 | 1.48 | 0.967 | 0.0721 |
| **B** | 5' | **1.69** | 0.937 | 0.04 | **1.54** | 0.956 | 0.0751 | **1.7** | 0.943 | 0.0189 | **1.47** | 0.967 | 0.0804 |

Table A7: **RIBO-former performances for different model optimization strategies and data sets**. Scores are calculated on the test set after selection of the model with the minimum validation loss (See Supplementary Figure A15, A16). For each data set and strategy, the cross-entropy loss ($\times 10^3$), area under the receiver operating characteristic curve (ROC), and area under the precision-recall curve (PR) are given. Results show the gain by having a model trained on a large variety of ribosome-profiling data sets using a supervised learning objective (Supplementary Table A3. All settings are evaluated using the same model architecture (Architecture 4, see Supplementary Table A5) and input token strategy (Strategy B, see Supplementary Table A6. The data is split in two folds (F1 and F2), with different parts of the transcriptome covered as training/validation/test data in each fold.

| | | **SRR1802129** | | | **SRR2433794** | | | **SRR2732970** | | | **SRR2733100** | | |
|---|---|---|---|---|---|---|---|---|---|---|---|---|---|
| | **Pre-train** | **Loss** | **ROC** | **PR** | **Loss** | **ROC** | **PR** | **Loss** | **ROC** | **PR** | **Loss** | **ROC** | **PR** |
| | - | 1.58 | 0.944 | 0.017 | 1.36 | 0.968 | 0.097 | 1.24 | 0.969 | 0.193 | 1.22 | 0.969 | 0.203 |
| **F1** | Supervised | **1.56** | 0.948 | 0.024 | **1.32** | 0.970 | 0.120 | **1.18** | 0.972 | 0.239 | **1.18** | 0.972 | 0.240 |
| | Self-Supervised | 1.57 | 0.946 | 0.020 | 1.36 | 0.966 | 0.110 | 1.20 | 0.970 | 0.227 | 1.20 | 0.969 | 0.226 |
| | - | 1.66 | 0.944 | 0.015 | 1.47 | 0.962 | 0.084 | 1.29 | 0.968 | 0.193 | 1.29 | 0.969 | 0.195 |
| **F2** | Supervised | **1.63** | 0.948 | 0.024 | **1.42** | 0.967 | 0.104 | **1.25** | 0.972 | 0.227 | **1.24** | 0.972 | 0.229 |
| | Self-Supervised | 1.65 | 0.945 | 0.021 | 1.44 | 0.965 | 0.098 | 1.26 | 0.969 | 0.214 | 1.28 | 0.969 | 0.214 |

| | | **SRR2954800** | | | **SRR8449577** | | | **SRR9113067** | | | **SRR11005875** | | |
|---|---|---|---|---|---|---|---|---|---|---|---|---|---|
| | **Position** | **Loss** | **ROC** | **PR** | **Loss** | **ROC** | **PR** | **Loss** | **ROC** | **PR** | **Loss** | **ROC** | **PR** |
| | - | 1.60 | 0.935 | 0.033 | 1.43 | 0.959 | 0.079 | 1.59 | 0.946 | 0.013 | 1.37 | 0.969 | 0.075 |
| **F1** | Supervised | **1.57** | 0.939 | 0.045 | **1.40** | 0.962 | 0.096 | **1.55** | 0.951 | 0.024 | **1.34** | 0.972 | 0.094 |
| | Self-Supervised | 1.59 | 0.936 | 0.036 | 1.42 | 0.959 | 0.087 | 1.58 | 0.946 | 0.019 | 1.37 | 0.968 | 0.077 |
| | - | 1.70 | 0.932 | 0.025 | 1.52 | 0.956 | 0.071 | 1.68 | 0.941 | 0.013 | 1.45 | 0.966 | 0.076 |
| **F2** | Supervised | **1.65** | 0.939 | 0.044 | **1.47** | 0.960 | 0.092 | **1.62** | 0.949 | 0.027 | **1.42** | 0.970 | 0.090 |
| | Self-Supervised | 1.68 | 0.933 | 0.033 | 1.50 | 0.958 | 0.085 | 1.67 | 0.944 | 0.015 | 1.45 | 0.967 | 0.079 |

Table A8: **Comparative performances of RIBO-former based on different subsets of the data**. The model predictions to detect translation initiation sites for each position on the trancriptome can be subsetted as a post-processing step. Using Ensembl translation initiation sites to derive a positive set, the area under the receiver operating characteristic curve (ROC) and area under the precision-recall curve (PR) are calculated. Given is the performance for all positions (total of ∼430M), with no conditions for what a valid ORF constitutes, positions that result in an ORF with a valid stop codon on the transcript (stop codon), an ORF length larger than 30 nucleotides (ORF length), and an ATG start codon (ATG start). Additionally, a subset has been selected using a minimum of 20 mapped reads (# Reads) on the transcript as a requirement. The performance and percentage of the total samples when using a combination of all listed conditions (Combined) is listed in the last set of columns. Note that the predictions of RIBO-former are those of the models pre-trained using a supervised learning strategy (see Supplementary Table 4), where the predictions of both models/folds are simply merged to cover the full transcriptome.

| | - | | Stop codon | | ORF length | | # Reads | | ATG start | | Combined | | |
|---|---|---|---|---|---|---|---|---|---|---|---|---|---|
| **Data set** | **ROC** | **PR** | **ROC** | **PR** | **ROC** | **PR** | **ROC** | **PR** | **ROC** | **PR** | **%** | **ROC** | **PR** |
| SRR1802129 | 0.945 | 0.020 | 0.946 | 0.020 | 0.943 | 0.021 | 0.981 | 0.041 | 0.952 | 0.318 | 0.6 | 0.983 | 0.502 |
| SRR2433794 | 0.965 | 0.101 | 0.966 | 0.103 | 0.964 | 0.103 | 0.983 | 0.137 | 0.966 | 0.419 | 1.0 | 0.981 | 0.516 |
| SRR2732970 | 0.969 | 0.220 | 0.970 | 0.222 | 0.968 | 0.223 | 0.986 | 0.285 | 0.958 | 0.399 | 1.1 | 0.972 | 0.487 |
| SRR2733100 | 0.969 | 0.215 | 0.970 | 0.217 | 0.968 | 0.218 | 0.986 | 0.279 | 0.957 | 0.399 | 1.1 | 0.971 | 0.488 |
| SRR2954800 | 0.935 | 0.034 | 0.935 | 0.034 | 0.932 | 0.036 | 0.972 | 0.068 | 0.942 | 0.266 | 0.6 | 0.974 | 0.417 |
| SRR8449577 | 0.958 | 0.083 | 0.959 | 0.084 | 0.957 | 0.085 | 0.983 | 0.128 | 0.961 | 0.382 | 0.8 | 0.982 | 0.514 |
| SRR9113067 | 0.944 | 0.016 | 0.945 | 0.016 | 0.942 | 0.017 | 0.971 | 0.024 | 0.950 | 0.289 | 0.9 | 0.974 | 0.399 |
| SRR11005875 | 0.967 | 0.077 | 0.968 | 0.078 | 0.966 | 0.080 | 0.985 | 0.106 | 0.969 | 0.432 | 1.0 | 0.984 | 0.534 |

Table A9: **Comparative performances of RIBO-former with different tools for *de novo* detection of expressed coding sequences**. Performances are given for multiple data sets and tools. Unlike RIBO-former, previous tools only evaluate a small selection of positions/ORFs on the transcriptome based on a variety of filters (e.g. start codon, ORF length, ...). Only these sites can be taken into account when calculating the score metric. Using annotated Ensembl coding sequenes (CDSs) that function as the positive set, the area under the receiver operating characteristic curve (ROC) and area under the precision-recall curve (PR) can be calculated. Some positions were left out as a mismatch exists between the given properties of both tools. Note that the number of positions and composition of positive and negative samples evaluated by each tool is unique and influences the score metric. Thus, performance scores are not comparable between tools. Rp-Bp and riboTISH were applied without RNA-seq data. Note that the predictions of RIBO-former are those of the models pre-trained using a supervised learning strategy (see Supplementary Table 4), where the predictions of both models/folds are simply merged to cover the full transcriptome.

| | | | | | | RIBO-former | |
|---|---|---|---|---|---|---|---|
| **Data set** | **ORFs** | **Ensembl CDS** | **Mismatch** | **ROC** | **PR** | **ROC** | **PR** |
| **PRICE** | | | | | | | |
| SRR1802129 | 6,417 | 352 (5.5%) | 57 (0.9%) | 0.786 | 0.159 | 0.968 | 0.639 |
| SRR2433794 | 42,978 | 2,534 (5.9%) | 422 (1.0%) | 0.693 | 0.185 | 0.961 | 0.639 |
| SRR2732970 | 109,391 | 7,478 (6.8%) | 1,178 (1.1%) | 0.635 | 0.276 | 0.965 | 0.720 |
| SRR2733100 | 122,202 | 4,735 (3.9%) | 1,112 (0.9%) | 0.562 | 0.153 | 0.977 | 0.694 |
| SRR2954800 | 7,654 | 637 (8.3%) | 194 (2.5%) | 0.726 | 0.170 | 0.935 | 0.604 |
| SRR8449577 | 15,520 | 1,027 (6.6%) | 267 (1.7%) | 0.728 | 0.207 | 0.958 | 0.650 |
| SRR9113067 | 11,944 | 457 (3.8%) | 193 (1.6%) | 0.781 | 0.195 | 0.967 | 0.565 |
| SRR11005875 | 13,175 | 1,477 (11.2%) | 141 (1.1%) | 0.721 | 0.246 | 0.943 | 0.678 |
| **Rp-Bp** | | | | | | | |
| SRR1802129 | 269,574 | 28,401 (10.5%) | 1,760 (0.7%) | 0.579 | 0.143 | 0.928 | 0.578 |
| SRR2433794 | 453,936 | 40,037 (8.8%) | 3,158 (0.7%) | 0.564 | 0.108 | 0.945 | 0.621 |
| SRR2732970 | 485,714 | 40,407 (8.3%) | 3,288 (0.7%) | 0.622 | 0.118 | 0.921 | 0.583 |
| SRR2733100 | 472,020 | 39,761 (8.4%) | 3,276 (0.7%) | 0.611 | 0.115 | 0.924 | 0.593 |
| SRR2954800 | 237,475 | 23,265 (9.8%) | 1,434 (0.6%) | 0.536 | 0.112 | 0.922 | 0.556 |
| SRR8449577 | 369,043 | 34,404 (9.3%) | 2,768 (0.8%) | 0.578 | 0.125 | 0.946 | 0.622 |
| SRR9113067 | 372,811 | 34,991 (9.4%) | 3,445 (0.9%) | 0.547 | 0.109 | 0.917 | 0.507 |
| SRR11005875 | 446,978 | 41,421 (9.3%) | 2,936 (0.7%) | 0.566 | 0.113 | 0.947 | 0.627 |
| **RiboCode** | | | | | | | |
| SRR1802129 | 60,026 | 21,510 (35.8%) | 6 (0.0%) | 0.657 | 0.523 | 0.790 | 0.642 |
| SRR2433794 | 134,404 | 46,001 (34.2%) | 19 (0.0%) | 0.756 | 0.603 | 0.806 | 0.651 |
| SRR2732970 | 145,781 | 42,850 (29.4%) | 28 (0.0%) | 0.784 | 0.575 | 0.812 | 0.647 |
| SRR2733100 | 145,781 | 42,850 (29.4%) | 28 (0.0%) | 0.784 | 0.575 | 0.816 | 0.651 |
| SRR2954800 | 22,624 | 8,427 (37.2%) | 5 (0.0%) | 0.582 | 0.466 | 0.838 | 0.723 |
| SRR8449577 | 107,173 | 39,945 (37.3%) | 30 (0.0%) | 0.731 | 0.605 | 0.799 | 0.665 |
| SRR9113067 | 58,462 | 21,412 (36.6%) | 22 (0.0%) | 0.630 | 0.507 | 0.770 | 0.622 |
| SRR11005875 | 125,964 | 47,249 (37.5%) | 23 (0.0%) | 0.731 | 0.609 | 0.784 | 0.650 |
| **RiboTISH** | | | | | | | |
| SRR1802129 | 646,112 | 50,144 (7.8%) | 36 (0.0%) | 0.583 | 0.105 | 0.932 | 0.527 |
| SRR2433794 | 1,020,518 | 69,479 (6.8%) | 442 (0.0%) | 0.556 | 0.078 | 0.941 | 0.558 |
| SRR2732970 | 1,039,658 | 69,286 (6.7%) | 121 (0.0%) | 0.572 | 0.078 | 0.915 | 0.526 |
| SRR2733100 | 1,002,275 | 67,270 (6.7%) | 85 (0.0%) | 0.565 | 0.077 | 0.919 | 0.535 |
| SRR2954800 | 442,725 | 34,448 (7.8%) | 993 (0.2%) | 0.532 | 0.085 | 0.931 | 0.544 |
| SRR8449577 | 842,566 | 60,171 (7.1%) | 433 (0.1%) | 0.574 | 0.089 | 0.945 | 0.568 |
| SRR9113067 | 759,182 | 55,710 (7.3%) | 2,418 (0.3%) | 0.553 | 0.086 | 0.924 | 0.472 |
| SRR11005875 | 1,006,709 | 71,656 (7.1%) | 1,163 (0.1%) | 0.566 | 0.085 | 0.944 | 0.563 |
| **Ribotricer** | | | | | | | |
| SRR1802129 | 238,095 | 23,206 (9.7%) | 4 (0.0%) | 0.569 | 0.111 | 0.966 | 0.709 |
| SRR2433794 | 506,515 | 33,181 (6.6%) | 16 (0.0%) | 0.594 | 0.078 | 0.974 | 0.692 |
| SRR2732970 | 487,689 | 28,823 (5.9%) | 15 (0.0%) | 0.557 | 0.065 | 0.971 | 0.687 |
| SRR2733100 | 171,597 | 4,022 (2.3%) | 5 (0.0%) | 0.603 | 0.034 | 0.964 | 0.447 |
| SRR2954800 | 155,014 | 15,248 (9.8%) | 8 (0.0%) | 0.586 | 0.122 | 0.959 | 0.679 |
| SRR8449577 | 363,661 | 28,043 (7.7%) | 7 (0.0%) | 0.592 | 0.093 | 0.973 | 0.706 |
| SRR9113067 | 195,331 | 15,107 (7.7%) | 3 (0.0%) | 0.593 | 0.099 | 0.962 | 0.646 |
| SRR11005875 | 439,689 | 34,384 (7.8%) | 11 (0.0%) | 0.582 | 0.091 | 0.971 | 0.694 |

Table A10: **RIBO-former performances for merged data sets**. Multiple data sets are merged by combining the mapped read counts along a transcript. This can improve performances and help pinpoint translated coding sequences. For each set of data, the cross-entropy loss ($\times 10^3$), area under the receiver operating characteristic curve (ROC), and area under the precision-recall curve (PR) are given. Note that the predictions of RIBO-former are those of the previously best performing set-up, following the supervised learning strategy (see Supplementary Table 4.

| | **Metric** | **SRR8449577** | **SRR8449577**<br>**SRR8449578** | **SRR9113067** | **SRR9113067**<br>**SRR9113068**<br>**SRR9113069** |
|---|---|---|---|---|---|
| **F2** Supervised | Loss | 1.47 | 1.36 | 1.62 | 1.51 |
| | ROC | 0.960 | 0.965 | 0.949 | 0.962 |
| | PR | 0.092 | 0.172 | 0.027 | 0.056 |

### 5. Supplementary Figures

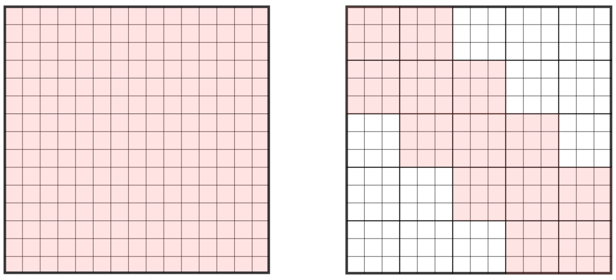

Figure A1: **Attention schemes used by the attention heads in the model. (left)** full attention heads allow every input nucleotide position to attend to all other positions on the transcript. These attention heads utilize the FAVOR+ approximation algorithm to allow input sequences of up to 30,000 tokens. **(right)**: Local attention is implemented by dividing the attention matrix in smaller blocks on which full attention is calculated. Three blocks around the evaluated input are calculated. The window size of each block listed as 'local_window_size' in Supplementary Table A5

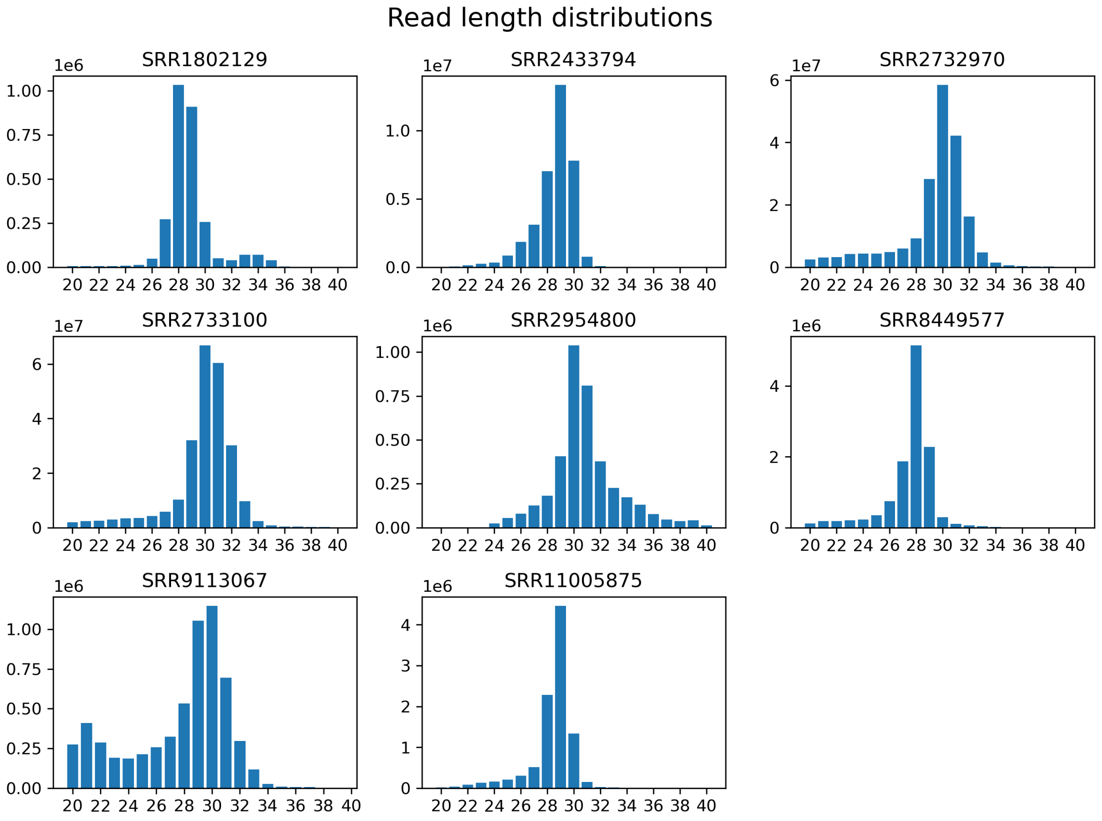


Figure A2: **Read length distributions for the evaluated datasets for reads mapped to the genome**. For each data set, the most abundant read length is generally around 29 nucleotides.

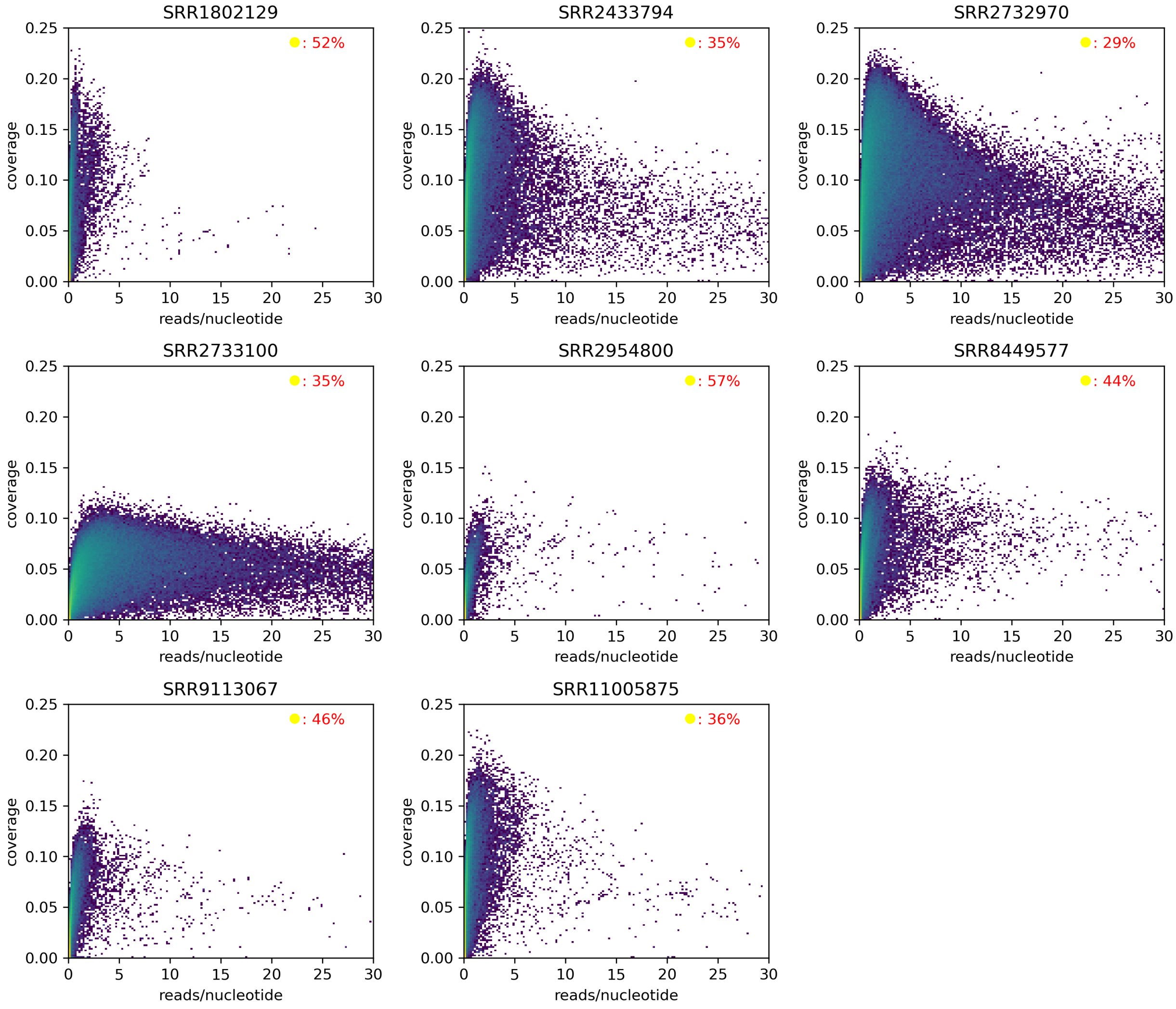


Figure A3: **2D histogram of transcript properties based on reads mapped.** For each data set, the percentage of transcripts in the transcriptome with practically no reads mapped (i.e., the bottom-left bin) is given (top-right corner) to reflect the expression profile of the tissue and/or quality of the data. The number of mapped reads are normalized by transcript length (reads/nucleotide). The coverage is calculated based on the percentage of the transcript positions that have at least one read mapped by their 5' position. The color map follows a logarithmic scale and is identical for all data sets.

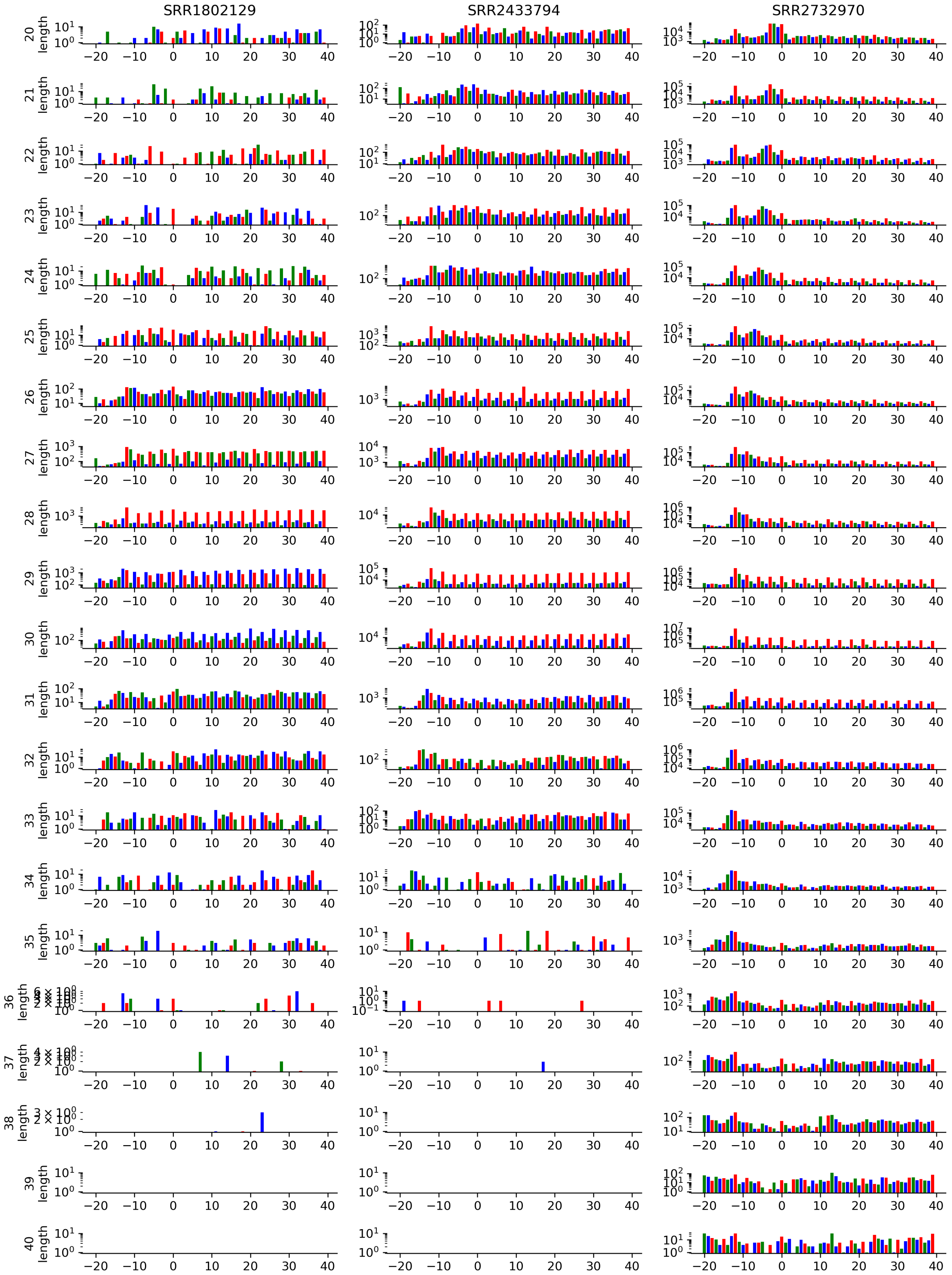


Figure A4: **Counts of reads mapped by their 5' positions along translation initiation start sites.** The figure showcases unique patterns of read alignments per read length and experiment. Read counts are taken by only evaluating translation initiation sites of coding sequences within the consensus coding sequence (CCDS) library. A window of 20 nucleotides upstream and 40 nucleotides downstream is taken. A logarithmic scale and alternating color scheme is used to highlight the patterns emerging from the triplet periodicity along the translation initiation site and coding sequence. Included are experiments SRR1802129, SRR2433794, and SRR2732970. Accompanied by Supplementary Figure A5 and A6.

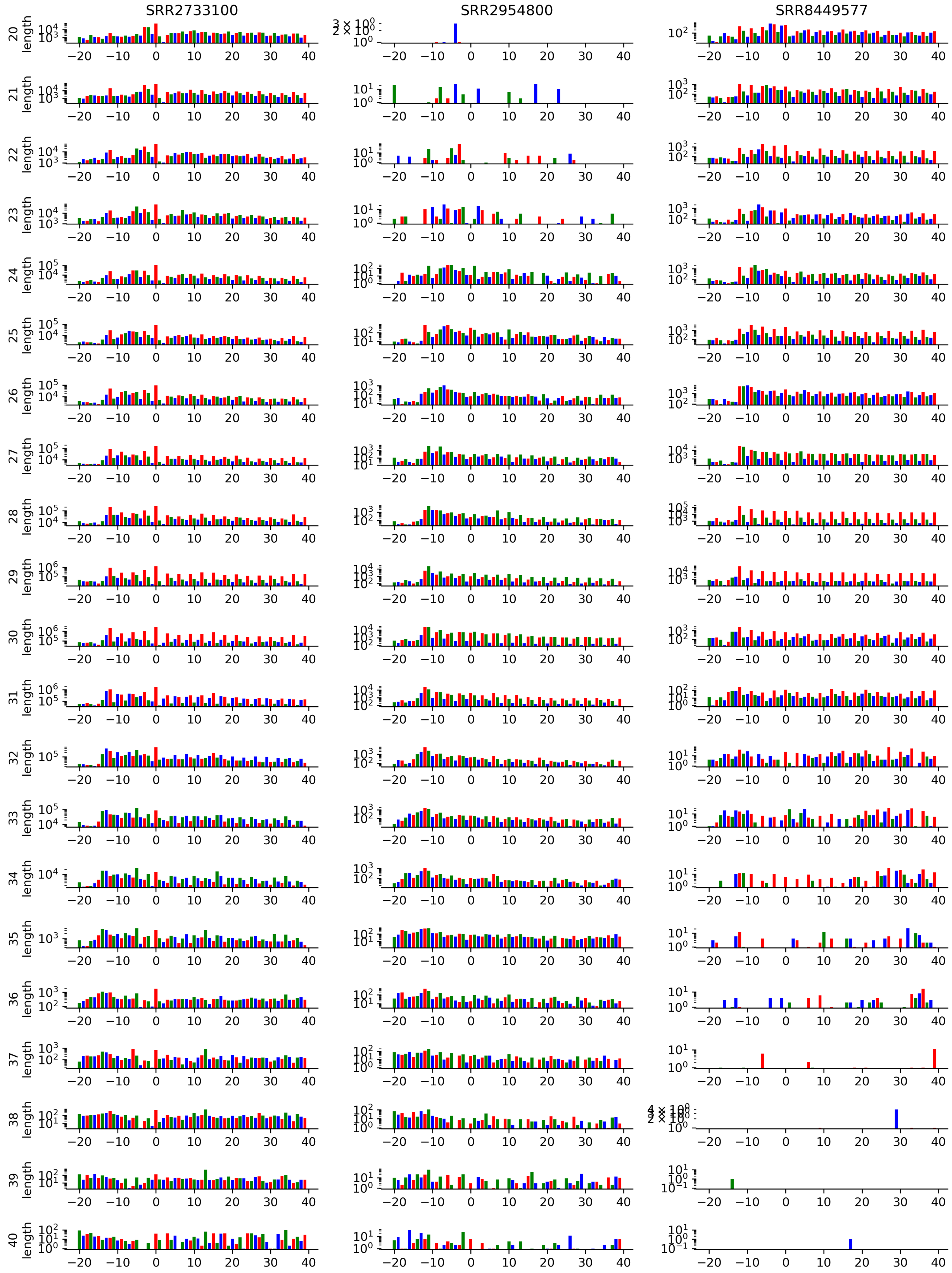


Figure A5: **Counts of reads mapped by their 5' positions along translation initiation start sites.** The figure showcases unique patterns of read alignments per read length and experiment. Read counts are taken by only evaluating translation initiation sites of coding sequences within the consensus coding sequence (CCDS) library. A window of 20 nucleotides upstream and 40 nucleotides downstream is taken. A logarithmic scale and alternating color scheme is used to highlight the patterns emerging from the triplet periodicity along the translation initiation site and coding sequence. Included are experiments SRR2733100, SRR2954800, and SRR8449577. Accompanied by Supplementary Figure A4 and A6.

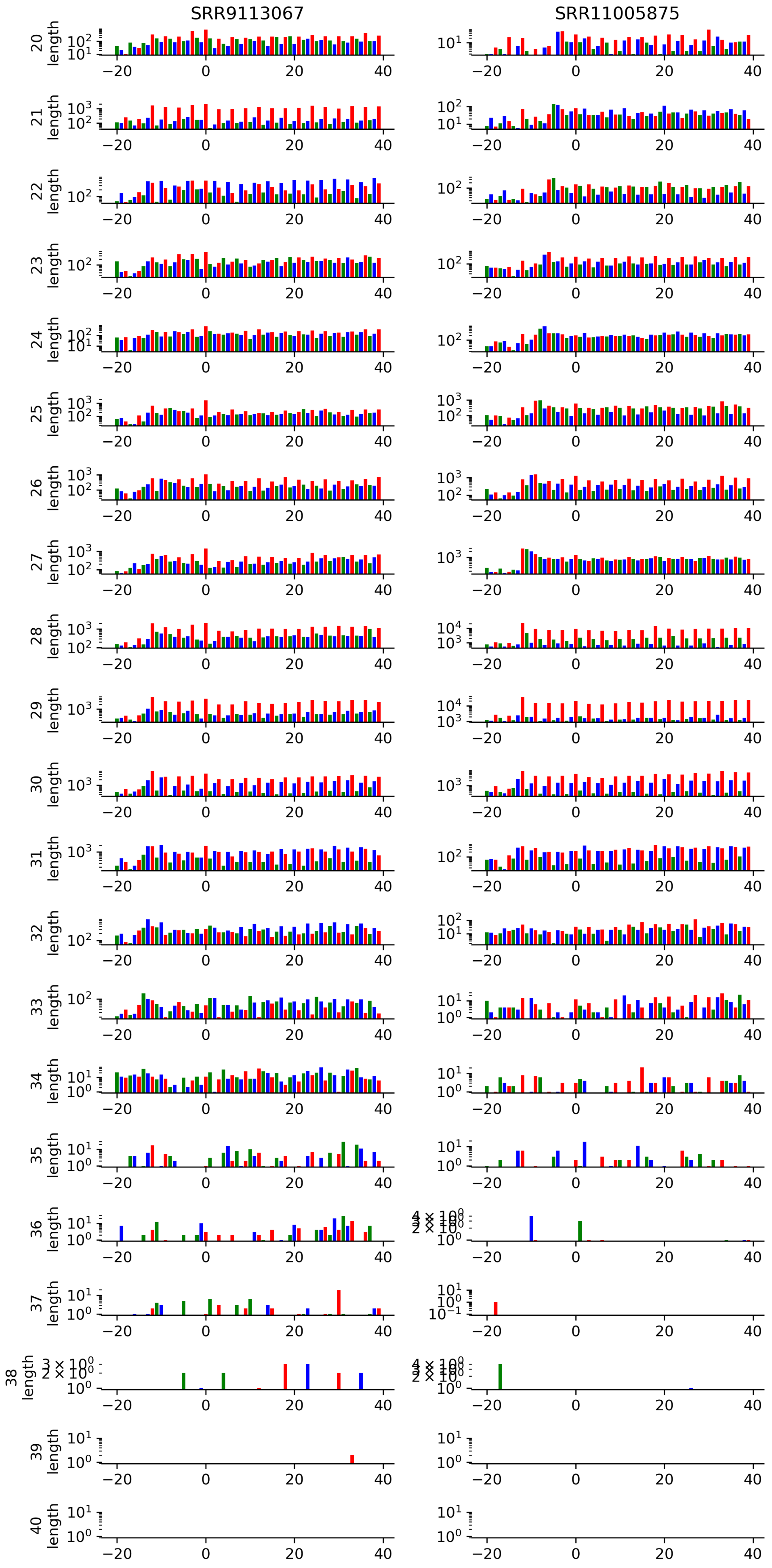


Figure A6: **Counts of reads mapped by their 5' positions along translation initiation start sites.** The figure showcases unique patterns of read alignments per read length and experiment. Read counts are taken by only evaluating translation initiation sites of coding sequences within the consensus coding sequence (CCDS) library. A window of 20 nucleotides upstream and 40 nucleotides downstream is taken. A logarithmic scale and alternating color scheme is used to highlight the patterns emerging from the triplet periodicity along the translation initiation site and coding sequence. Included are experiments SRR9113067, SRR11005875. Accompanied by Supplementary Figure A4 and A5.

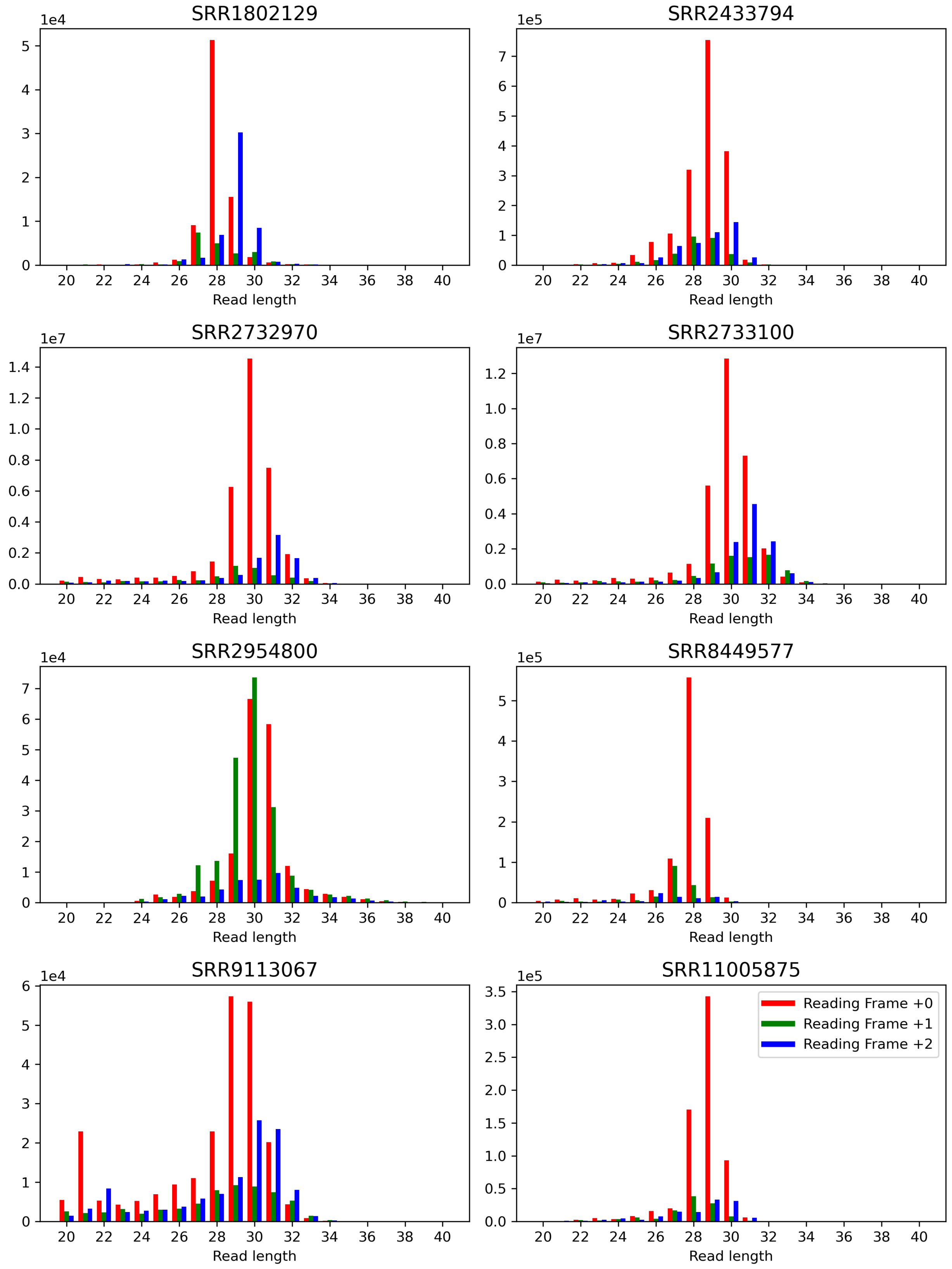


Figure A7: **Read length counts binned by reading frame offset.** Reads are mapped by their 5' positions. The figure highlights the skewed abundance of reads as influenced by the reading frame of the neighboring translation initiation site. Similar plots have been used to offset the mapping position of reads in relation to their length. Read counts are taken by only evaluating translation initiation sites of coding sequences within the consensus coding sequence (CCDS) library. A window of 20 nucleotides upstream and 40 nucleotides downstream is taken to calculate the total read counts.

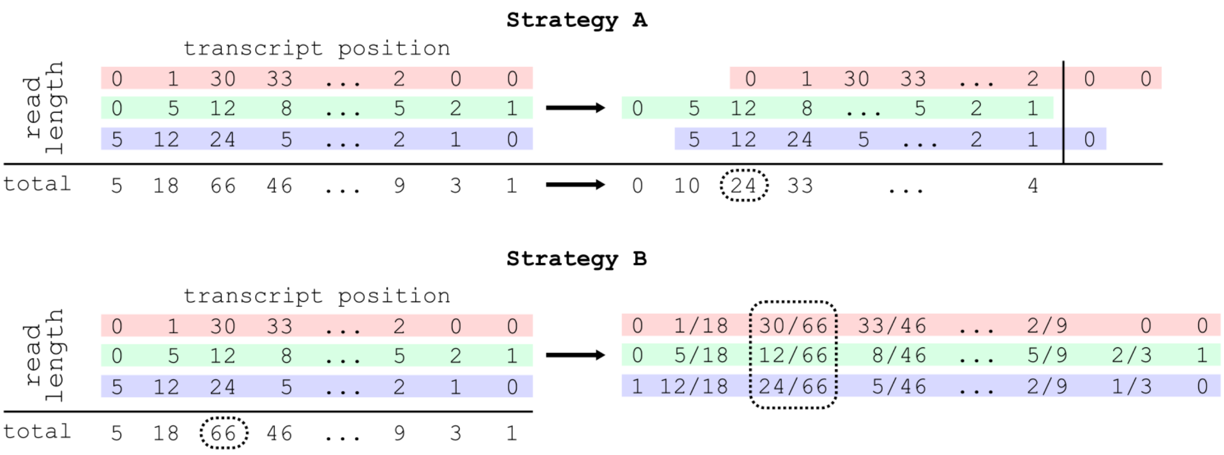


Figure A8: **Illustration of the data applied for calculating the input vector representation.** For a given matrix containing reads mapped according to their 5'-end by transcript position and read length. Strategy A: reads are offset according to a fixed value for each read length. The total read count is applied for further processing. Strategy B: both the total read count and the fractional abundance of each read length is used to obtain an input vector representation. Input vector representations are calculated for each position (e.g. dotted square encapsulates data used for a single position). Note that in contrast to the illustration, data is generally sparse and ribosome profiling data is applied for 21 read lengths ($[20, 40]$). See Supplementary Figure A9 on how the data of both strategies is used to create the input tokens.

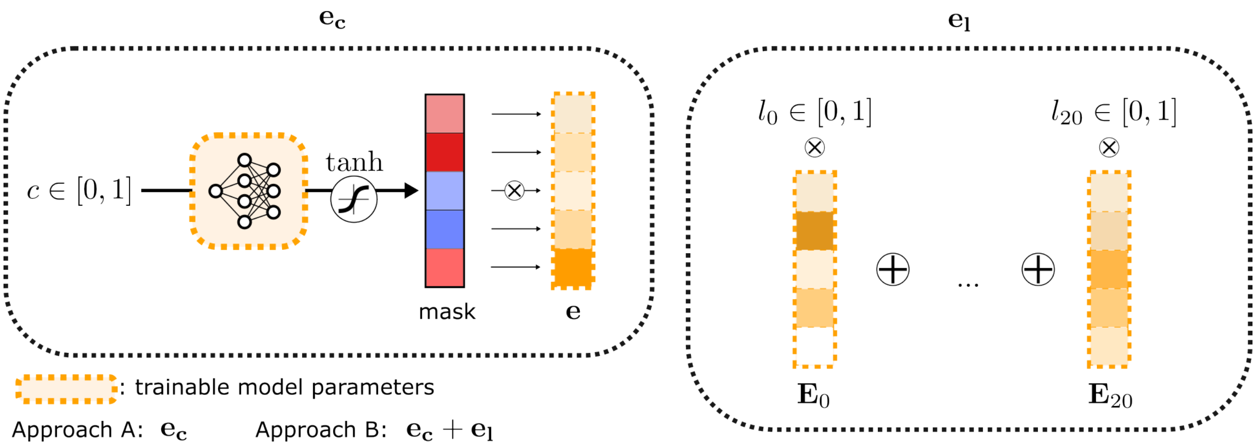


Figure A9: **Scheme of two different strategies for constructing the input vector.** See Supplementary Figure A8 on the inputs generated from mapped reads. Strategy A: the normalized read count is fed into short feed-forward neural network to obtain an input vector. Strategy B: A unique vector embedding is optimized as part of the model training process. For a given input, these embeddings are multiplied by the fractional representation of that read length at that position. The final vector is obtained through the summation of the weighted vector embeddings and count embedding. Read count $c \in [0, 1]$, read count embedding $\mathbf{e} \in \mathbb{R}^h$, read length embeddings $\mathbf{E} \in \mathbb{R}^{21 \times h}$ and read length fractions $\mathbf{l} \in [0, 1]^{21}$, where $\sum \mathbf{l} = 1$. The matrix $\mathbf{E}$ incorporates vector embeddings for read lengths 20–40 and is optimized as part of the training process. Note that data is generally sparse and the summation of 22 vectors for strategy B constitute mostly zero vectors.

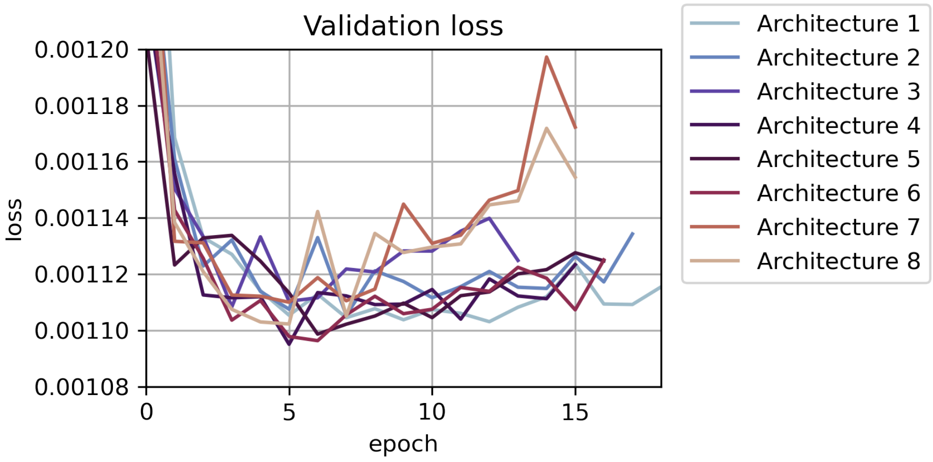


Figure A10: **The loss curves of the model architectures trained for detecting TIS using ribosome profiling data.** The validation sets used are chromosomes 2 and 14. The hyperparameters for each model are given in Table A5. Architectures 1–2 converge slowly over several epochs without reaching a minimum within the evaluated time frame, indicating too few model weights. The higher number of model weights of architectures 7–8 result in clear overfitting from epoch 5 onward. Architecture 4 returns the lowest loss, and is selected for model benchmarking. While the minimum loss is similar for all architectures, the plot confirms our selection of a model architecture with a suitable number of parameters.

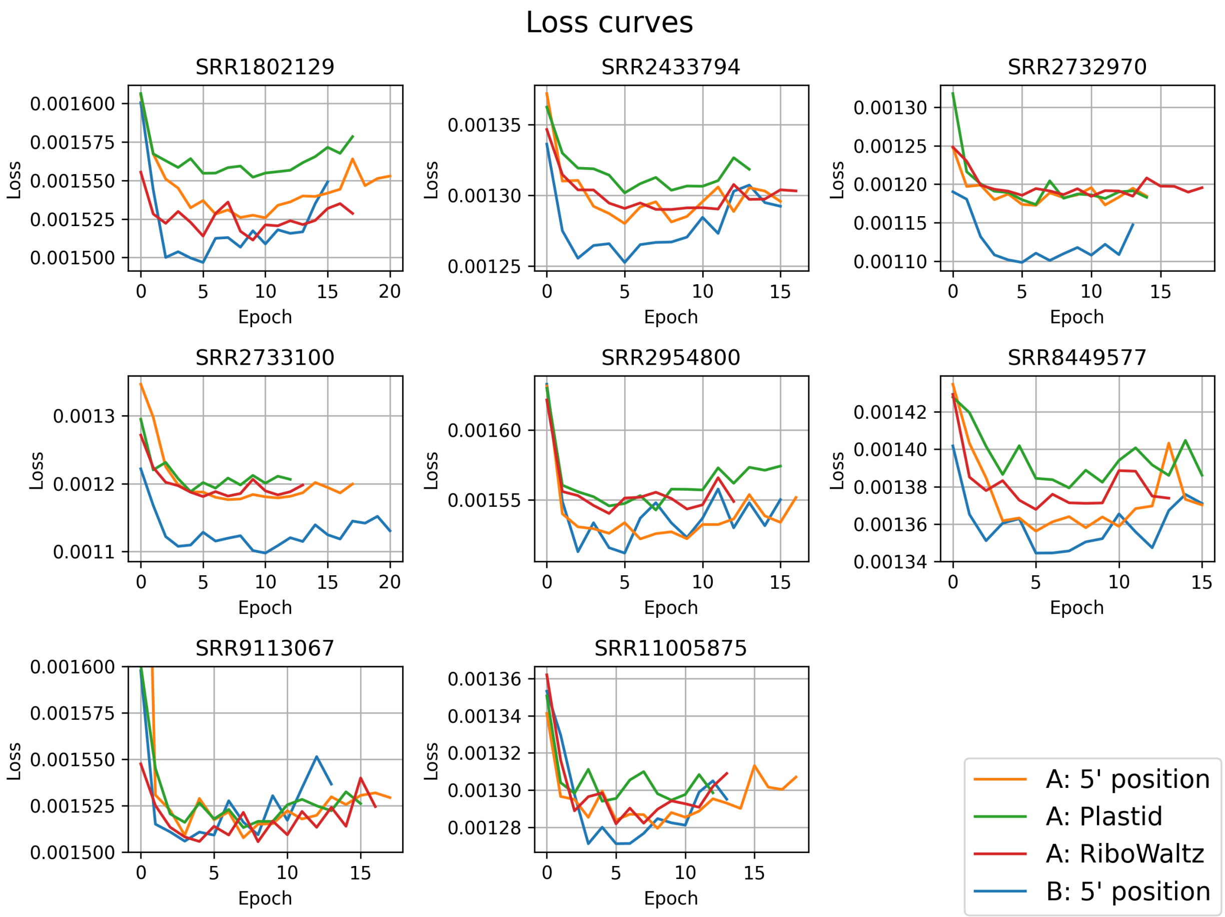


Figure A11: **Validation loss for different input token strategies and data sets throughout the training process.** Results indicate the relevance of read length information for the prediction of translation initiation sites using ribosome profiling data, especially for data sets featuring a higher read depth (see Supplementary Table A3). All strategies are evaluated using the same model architecture and training/validation data (Architecture 4, see Supplementary Table A5, Supplementary Figure A10), Strategy A generates input tokens for the model utilizing read count information for every position of the transcript. Strategy A includes mappings generated by taking the 5' position of every read, and offsetting reads based on read length following two strategies (Plastid, RiboWaltz). Strategy B includes information on both the positions and read lengths of the mapped reads.

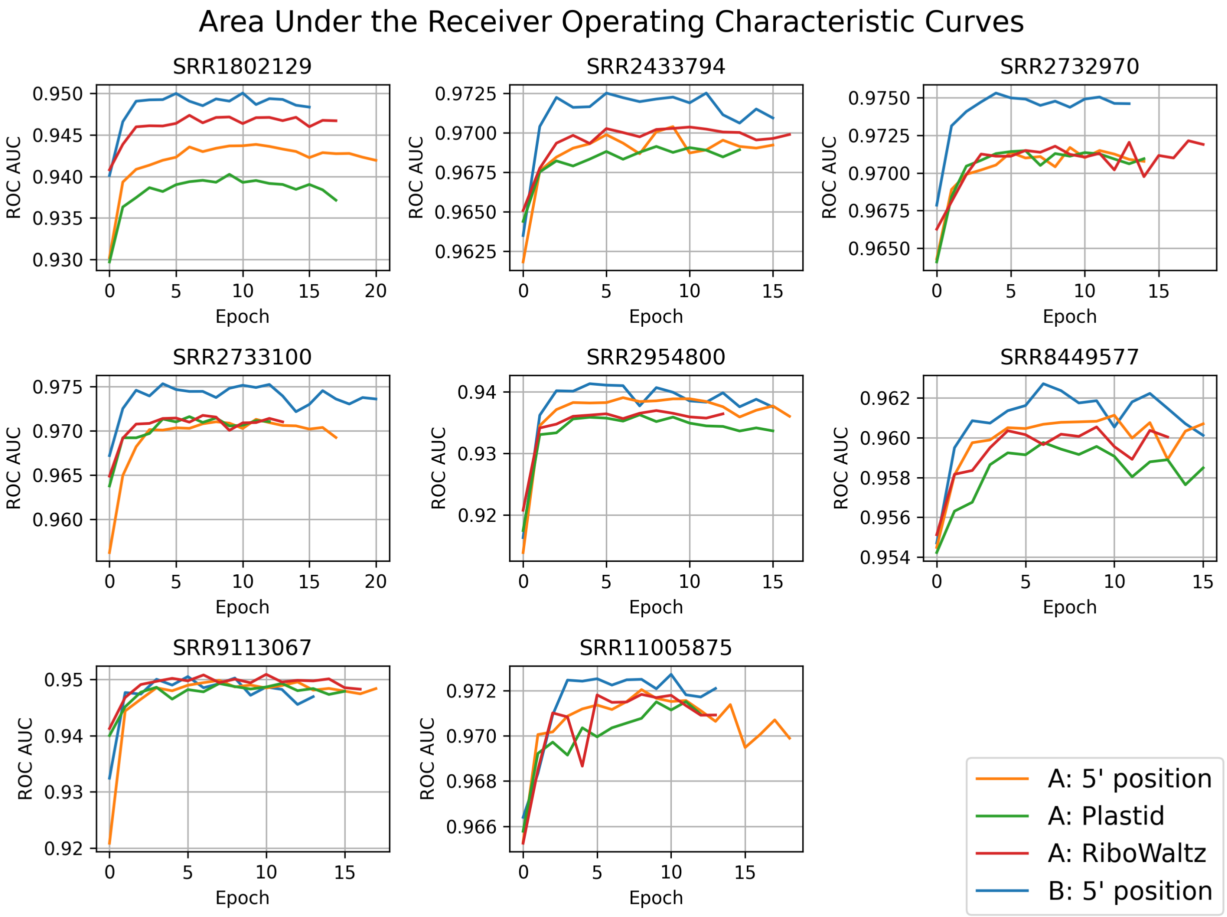


Figure A12: **Area under the receiver operating characteristic curve (ROC AUC) scores for different input token strategies and data sets.** Scores are given for every epoch throughout the training process. Results indicate the relevance of read length information for the prediction of translation initiation sites using ribosome profiling data, especially for data sets featuring a higher read depth (see Supplementary Table A3). All strategies are evaluated using the same model architecture and training/validation data (Architecture 4, see Supplementary Table A5, Supplementary Figure A10), Strategy A generates input tokens for the model utilizing read count information for every position of the transcript. Strategy A includes mappings generated by taking the 5' position of every read, and offsetting reads based on read length following two strategies (Plastid, RiboWaltz). Strategy B includes information on both the positions and read lengths of the mapped reads.

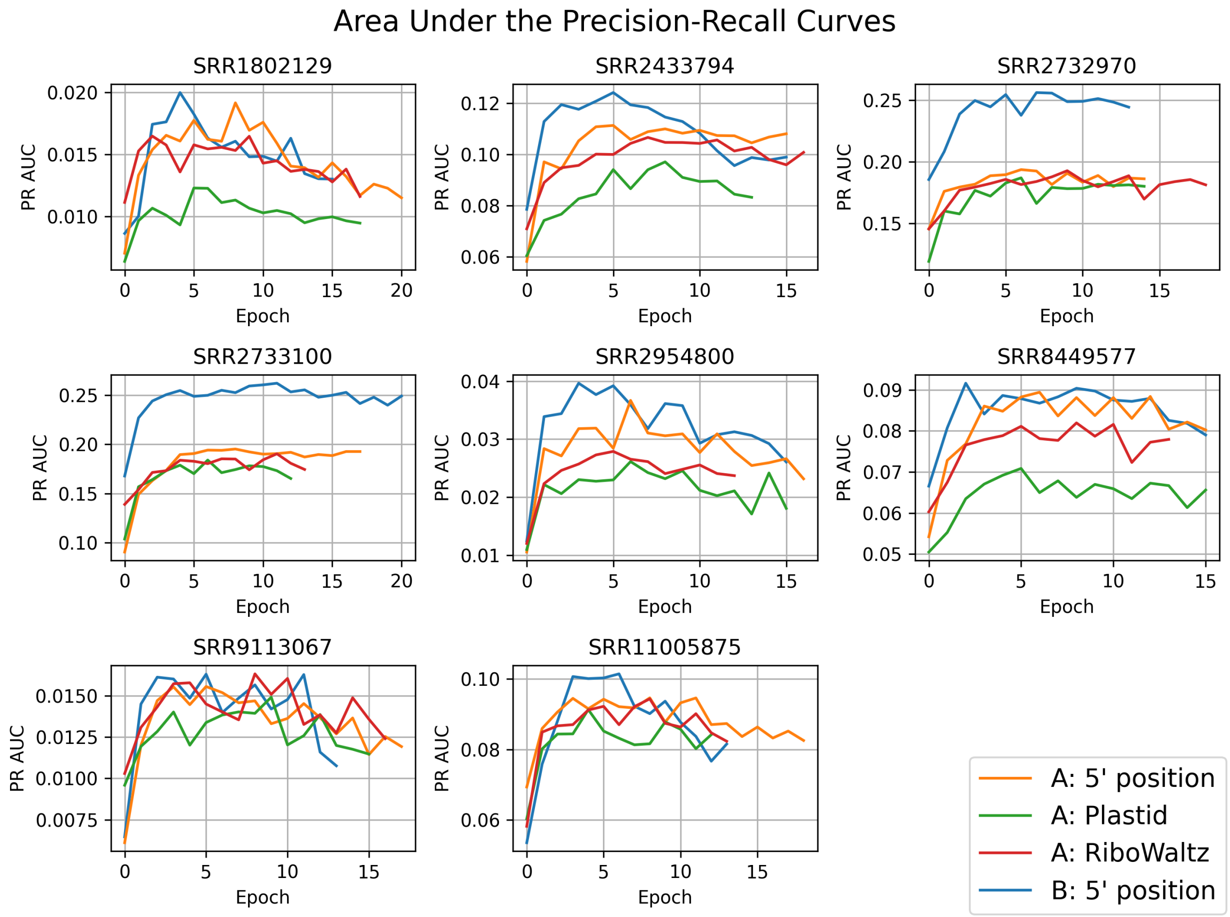


Figure A13: **Area under the precision-recall scores for different input token strategies and data sets.** Scores are given for every epoch throughout the training process. Results indicate the relevance of read length information for the prediction of translation initiation sites using ribosome profiling data, especially for data sets featuring a higher read depth (see Supplementary Table A3). All strategies are evaluated using the same model architecture and training/validation data (Architecture 4, see Supplementary Table A5, Supplementary Figure A10), Strategy A generates input tokens for the model utilizing read count information for every position of the transcript. Strategy A includes mappings generated by taking the 5' position of every read, and offsetting reads based on read length following two strategies (Plastid, RiboWaltz). Strategy B includes information on both the positions and read lengths of the mapped reads.

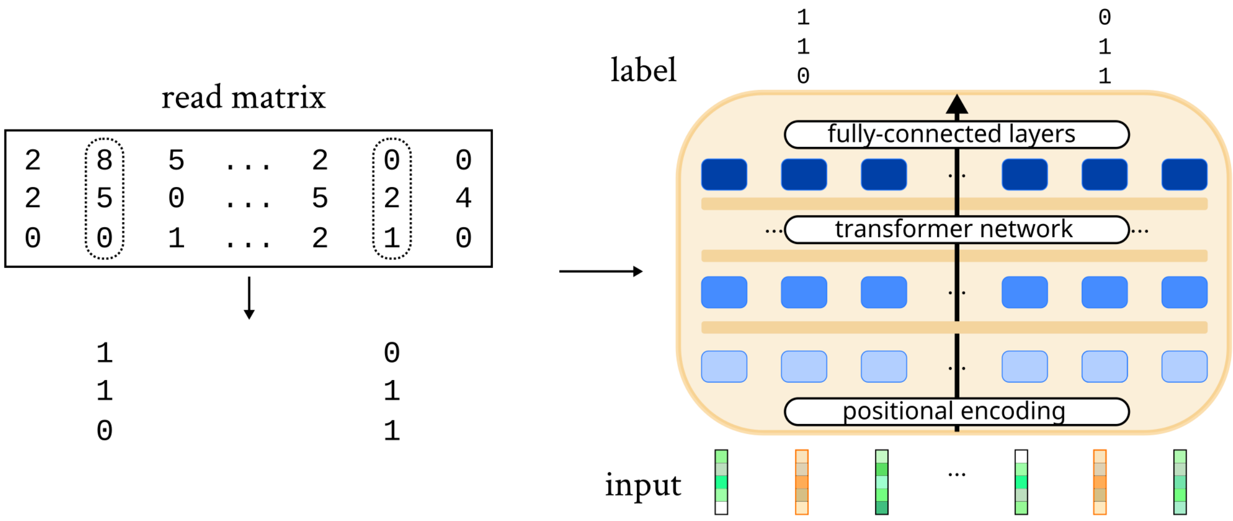


Figure A14: **Self-supervised learning implementation for ribosome profiling data.** One of the pre-training approaches investigated in this paper. A model is trained to infer the presence of a mapped ribosome reads at a given position. The task constitutes a binary multi-label classification task. 15% of the input positions were randomly selected (dotted frame) and masked using a custom input embedding (orange input vectors). Positive labels are allocated to read lengths having more than one read mapped at a given position. Note that in contrast to the illustration, data is generally sparse and ribosome profiling data is applied for 21 read lengths ($[20, 40]$).

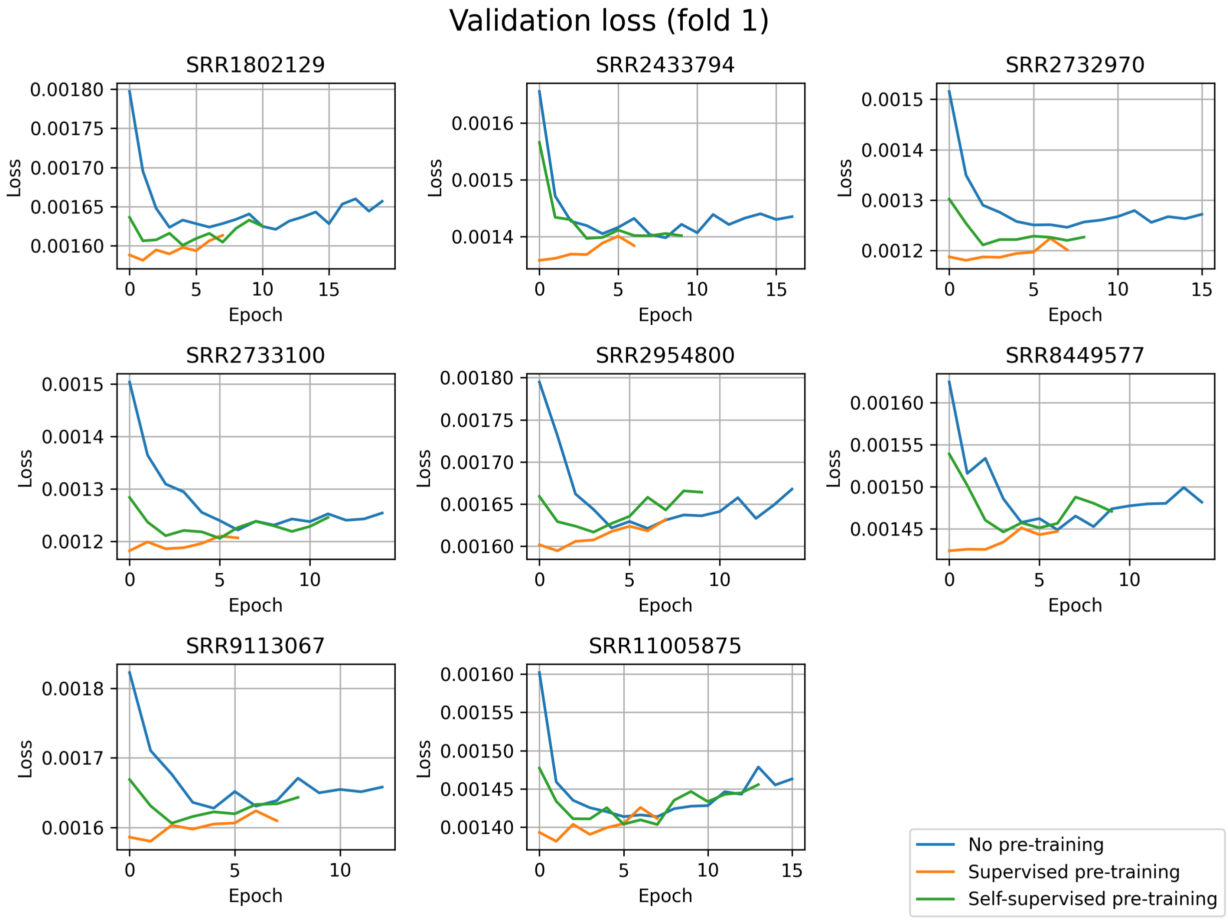


Figure A15: **Validation cross-entropy loss of different training schemes of RIBO-former.** Eight data sets were evaluated following three approaches. This is achieved by training a model from scratch (no pre-training) or using a pre-trained model fit on a selection of eight separate data sets (see Supplementary Table A3). Pre-trained models include both those fit following a supervised learning objective (supervised pre-training) on identifying translation initiation sites, and those fit following a self-supervised learning objective, similar to those found in language processing (see Supplementary Figure A14). This figure shows the models trained on chromosomes 3, 5, 7, 11, 13, 15, 19, 21, and X with chromosomes 1, 9, and 17 used as validation set.

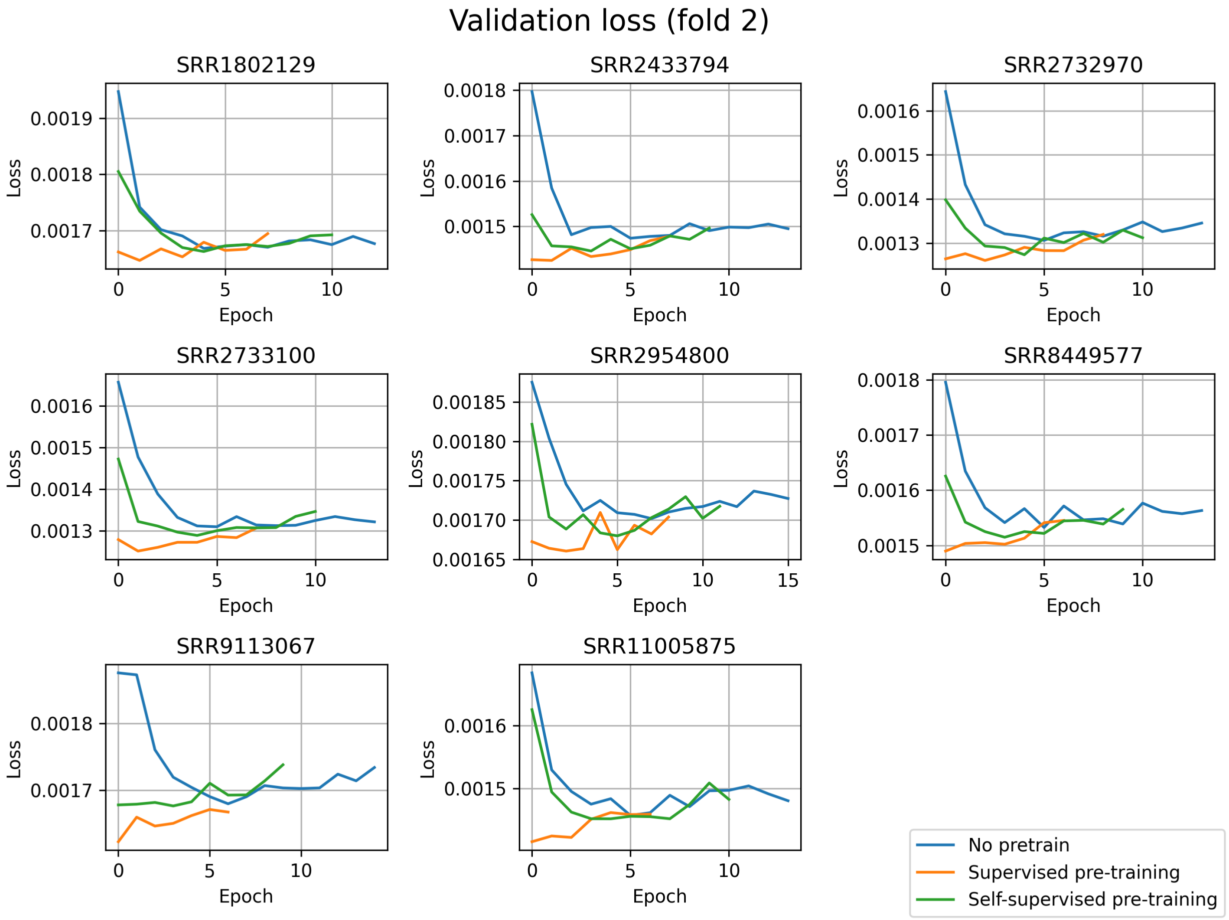


Figure A16: **Validation cross-entropy loss of different training schemes of RIBO-former.** Eight data sets were evaluated following three approaches. This is achieved by training a model from scratch (no pre-training) or using a pre-trained model fit on a selection of eight separate data sets (see Supplementary Table A3). Pre-trained models include both those fit following a supervised learning objective (supervised pre-training) on identifying translation initiation sites, and those fit following a self-supervised learning objective, similar to those found in language processing (see Supplementary Figure A14). This figure shows the models trained on chromosomes 2, 6, 8, 10, 14, 16, 18, 22, and Y with chromosomes 4, 12, and 20 used as validation set.

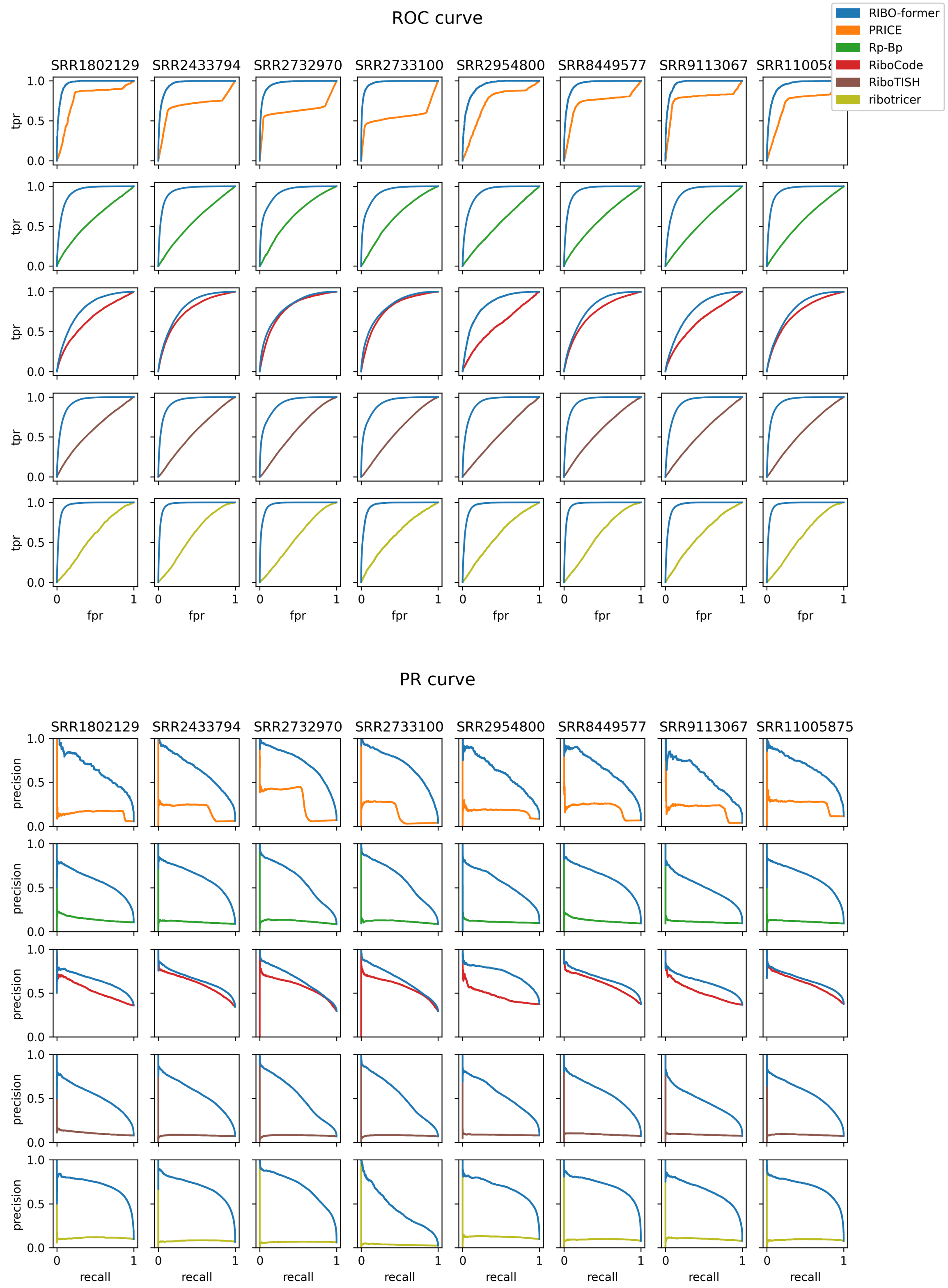


Figure A17: **Receiver operating characteristics and Precision-recall curves for each of the evaluated data sets.** Performances are given for multiple data sets and tools. Unlike RIBO-former, previous tools only evaluate a small selection of positions/ORFs on the transcriptome based on a variety of filters (e.g. start codon, ORF length, ...). Only these sites can be taken into account when calculating the score metric. Using annotated Ensembl coding sequenes (CDSs) that function as the positive set, the receiver operating characteristic curve (ROC) and precision-recall curve (PR) are calculated. the area under the receiver operating characteristic curve (ROC) and area under the precision-recall curve (PR) can be calculated. Rp-Bp and riboTISH were applied without RNA-seq data. Note that the number of positions and composition of positive and negative samples evaluated by each tool is unique and influences the score metric. Thus, performance scores are not comparable between tools. Supplementary Table A9 gives a more complete and accurate overview of the benchmark listed by tool and data set, including metadata on the number of positions evaluated and size of the positive set.

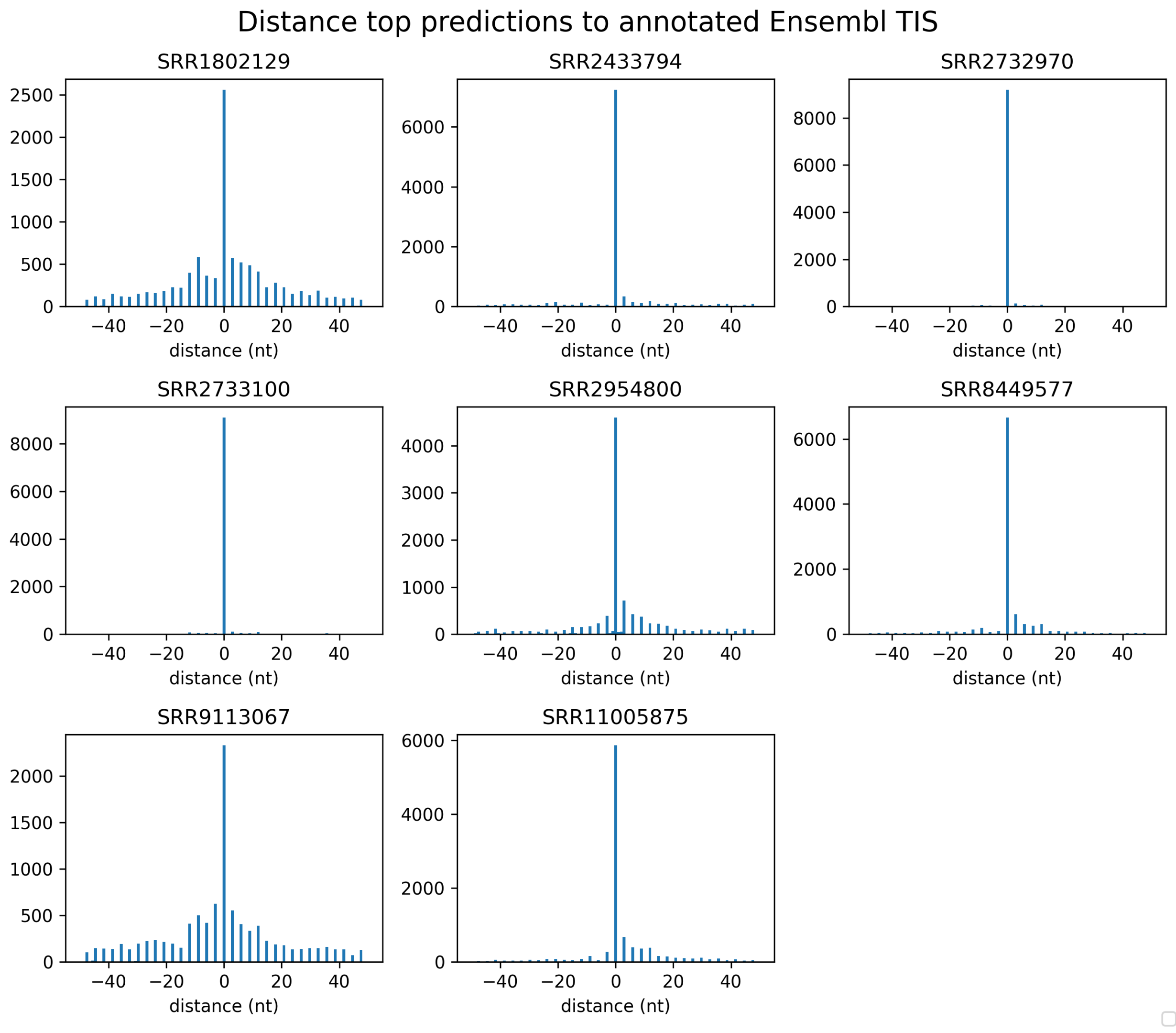


Figure A18: **Accuracy of pinpointing translation initiation site (TIS) positions is correlated with read depth of ribosome profiling experiment.** Because RIBO-former does not pre-process candidate open reading frames or pass start codon information, pinpointing TISs can suffer for data sets with lower read depth. For each data set, the histograms on distances between the a top-scoring RIBO-former prediction and an annotated Ensembl TIS is given. Interestingly, the figure proves RIBO-former capable of determining the correct reading frame for all data sets.

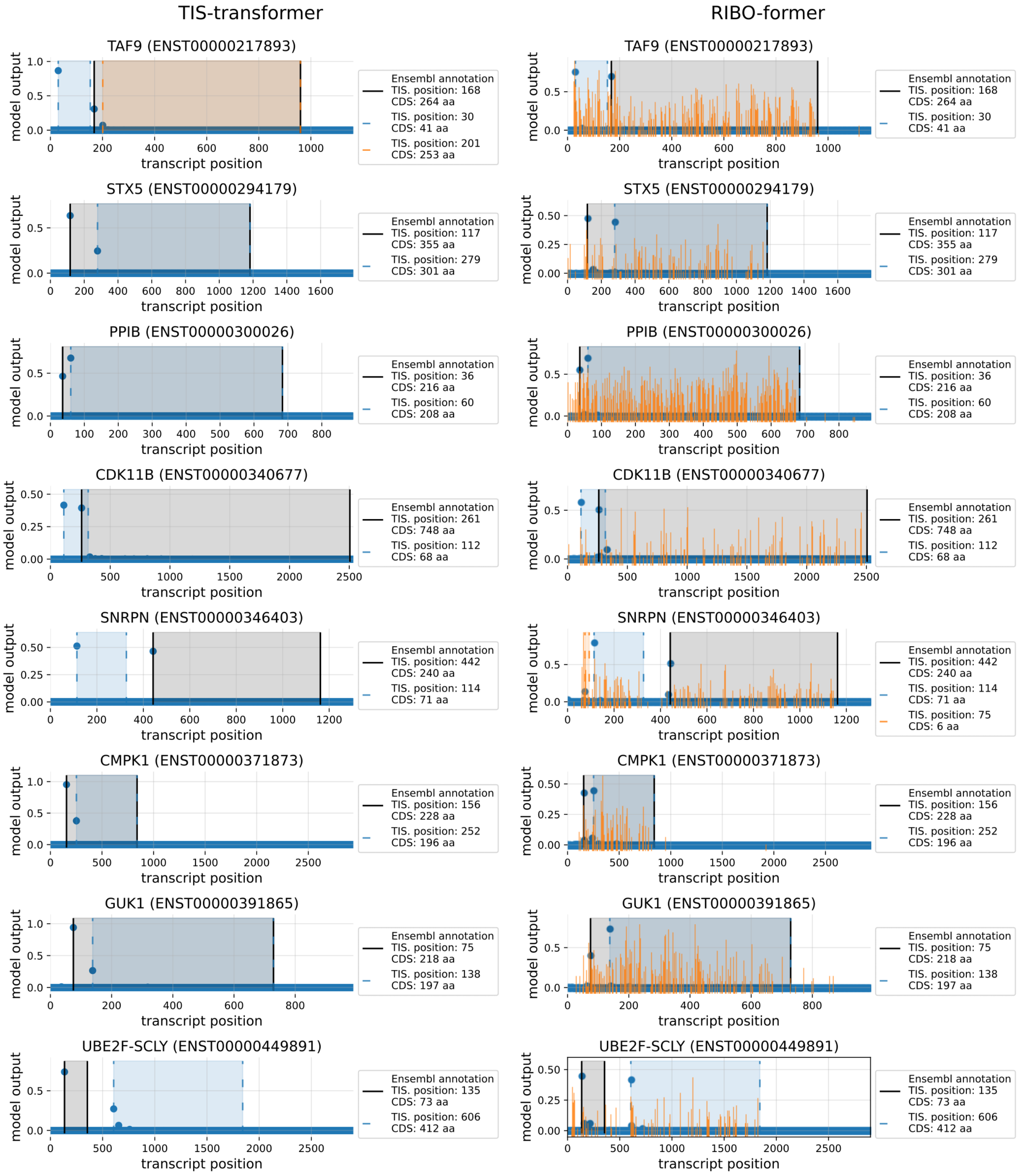


Figure A19: **Transcripts with multiple translation initiation sites are detected by both RIBO-former and TIS transformer.** Given are the model outputs (y-axis) for each position of the transcript (x-axis) for TIS transformer (left) and RIBO-former (rigth; data set SRR2732970). Mapped read counts used as input to RIBO-former are given in bright orange and displayed on a logarithmic scale. For high model predictions, the bounds of the resulting CDS are given, as well as their length. When a CDS is present in Ensembl, the bounds are represented by full black lines. Note that TIS transformer only processes transcript sequence information while RIBO-former only processes ribosome profiling information.

### References

N. Ahmed, P. Sormanni, P. Ciryam, M. Vendruscolo, C. M. Dobson, and E. P. O'Brien. Identifying A- and P-site locations on ribosome-protected mRNA fragments using Integer Programming. *Scientific Reports*, 9(1):6256, Apr. 2019. ISSN 2045-2322. doi: 10.1038/s41598-019-42348-x.

M. Bencun, O. Klinke, A. Hotz-Wagenblatt, S. Klaus, M.-H. Tsai, R. Poirey, and H.-J. Delecluse. Translational profiling of B cells infected with the Epstein-Barr virus reveals 5′ leader ribosome recruitment through upstream open reading frames. *Nucleic Acids Research*, 46(6):2802–2819, Apr. 2018. ISSN 0305-1048. doi: 10.1093/nar/gky129.

L. Calviello, N. Mukherjee, E. Wyler, H. Zauber, A. Hirsekorn, M. Selbach, M. Landthaler, B. Obermayer, and U. Ohler. Detecting actively translated open reading frames in ribosome profiling data. *Nature Methods*, 13(2):165–170, Feb. 2016. ISSN 1548-7105. doi: 10.1038/nmeth.3688.

J. Chen, A.-D. Brunner, J. Z. Cogan, J. K. Nuñez, A. P. Fields, B. Adamson, D. N. Itzhak, J. Y. Li, M. Mann, M. D. Leonetti, and J. S. Weissman. Pervasive functional translation of noncanonical human open reading frames. *Science*, 367(6482):1140–1146, Mar. 2020. doi: 10.1126/science.aay0262.

K. Choromanski, V. Likhosherstov, D. Dohan, X. Song, A. Gane, T. Sarlos, P. Hawkins, J. Davis, A. Mohiuddin, L. Kaiser, D. Belanger, L. Colwell, and A. Weller. Rethinking Attention with Performers. *arXiv:2009.14794 [cs, stat]*, Mar. 2021.

S. Choudhary, W. Li, and A. D. Smith. Accurate detection of short and long active ORFs using Ribo-seq data. *Bioinformatics*, 36(7):2053–2059, Apr. 2020. ISSN 1367-4803. doi: 10.1093/bioinformatics/btz878.

S. Y. Chun, C. M. Rodriguez, P. K. Todd, and R. E. Mills. SPECtre: A spectral coherence-based classifier of actively translated transcripts from ribosome profiling sequence data. *BMC Bioinformatics*, 17(1):482, Nov. 2016. ISSN 1471-2105. doi: 10.1186/s12859-016-1355-4.

J. Clauwaert, Z. McVey, R. Gupta, and G. Menschaert. TIS Transformer: Remapping the human proteome using deep learning. *NAR genomics and bioinformatics*, 5(1):lqad021, Mar. 2023. ISSN 2631-9268. doi: 10.1093/nargab/lqad021.

J. Crappé, E. Ndah, A. Koch, S. Steyaert, D. Gawron, S. De Keulenaer, E. De Meester, T. De Meyer, W. Van Criekinge, P. Van Damme, and G. Menschaert. PROTEOFORMER: Deep proteome coverage through ribosome profiling and MS integration. *Nucleic Acids Research*, 43(5):e29, Mar. 2015. ISSN 0305-1048. doi: 10.1093/nar/gku1283.

J. G. Dunn and J. S. Weissman. Plastid: nucleotide-resolution analysis of next-generation sequencing and genomics data. *BMC Genomics*, 17(1):958, 2016. ISSN 1471-2164. doi: 10.1186/s12864-016-3278-x. URL `http://dx.doi.org/10.1186/s12864-016-3278-x`.

F. Erhard, A. Halenius, C. Zimmermann, A. L'Hernault, D. J. Kowalewski, M. P. Weekes, S. Stevanovic, R. Zimmer, and L. Dölken. Improved Ribo-seq enables identification of cryptic translation events. *Nature Methods*, 15(5):363–366, May 2018. ISSN 1548-7105. doi: 10.1038/nmeth.4631.

H. Fang, Y.-F. Huang, A. Radhakrishnan, A. Siepel, G. J. Lyon, and M. C. Schatz. Scikit-ribo Enables Accurate Estimation and Robust Modeling of Translation Dynamics at Codon Resolution. *Cell Systems*, 6(2):180–191.e4, Feb. 2018. ISSN 2405-4712. doi: 10.1016/j.cels.2017.12.007.

A. P. Fields, E. H. Rodriguez, M. Jovanovic, N. Stern-Ginossar, B. J. Haas, P. Mertins, R. Raychowdhury, N. Hacohen, S. A. Carr, N. T. Ingolia, A. Regev, and J. S. Weissman. A Regression-Based Analysis of Ribosome-Profiling Data Reveals a Conserved Complexity to Mammalian Translation. *Molecular Cell*, 60 (5):816–827, Dec. 2015. ISSN 1097-2765. doi: 10.1016/j.molcel.2015.11.013.

B. Gaertner, S. van Heesch, V. Schneider-Lunitz, J. F. Schulz, F. Witte, S. Blachut, S. Nguyen, R. Wong, I. Matta, N. Hübner, and M. Sander. A human ESC-based screen identifies a role for the translated lncRNA LINC00261 in pancreatic endocrine differentiation. *eLife*, 9:e58659, Aug. 2020. ISSN 2050-084X. doi: 10.7554/eLife.58659.

D. Gawron, E. Ndah, K. Gevaert, and P. Van Damme. Positional proteomics reveals differences in N-terminal proteoform stability. *Molecular Systems Biology*, 12(2):858, Feb. 2016. ISSN 1744-4292. doi: 10.15252/msb.20156662.

C. Gonzalez, J. S. Sims, N. Hornstein, A. Mela, F. Garcia, L. Lei, D. A. Gass, B. Amendolara, J. N. Bruce, P. Canoll, and P. A. Sims. Ribosome Profiling Reveals a Cell-Type-Specific Translational Landscape in Brain Tumors. *Journal of Neuroscience*, 34(33):10924–10936, Aug. 2014. ISSN 0270-6474, 1529-2401. doi: 10.1523/JNEUROSCI.0084-14.2014.

Z. Ji. RibORF: Identifying Genome-Wide Translated Open Reading Frames Using Ribosome Profiling. *Current Protocols in Molecular Biology*, 124(1):e67, 2018. ISSN 1934-3647. doi: 10.1002/cpmb.67.

Z. Ji, R. Song, A. Regev, and K. Struhl. Many lncRNAs, 5'UTRs, and pseudogenes are translated and some are likely to express functional proteins. *eLife*, 4:e08890, Dec. 2015. ISSN 2050-084X. doi: 10.7554/eLife.08890.

F. Lauria, T. Tebaldi, P. Bernabò, E. J. N. Groen, T. H. Gillingwater, and G. Viero. riboWaltz: Optimization of ribosome P-site positioning in ribosome profiling data. *PLOS Computational Biology*, 14(8):e1006169, Aug. 2018. ISSN 1553-7358. doi: 10.1371/journal.pcbi.1006169.

F. Loayza-Puch, K. Rooijers, L. C. M. Buil, J. Zijlstra, J. F. Oude Vrielink, R. Lopes, A. P. Ugalde, P. van Breugel, I. Hofland, J. Wesseling, O. van Tellingen, A. Bex, and R. Agami. Tumour-specific proline vulnerability uncovered by differential ribosome codon reading. *Nature*, 530(7591):490–494, Feb. 2016. ISSN 1476-4687. doi: 10.1038/nature16982.

B. Malone, I. Atanassov, F. Aeschimann, X. Li, H. Großhans, and C. Dieterich. Bayesian prediction of RNA translation from ribosome profiling. *Nucleic Acids Research*, 45(6):2960–2972, Apr. 2017. ISSN 0305-1048. doi: 10.1093/nar/gkw1350.

T. F. Martinez, Q. Chu, C. Donaldson, D. Tan, M. N. Shokhirev, and A. Saghatelian. Accurate annotation of human protein-coding small open reading frames. *Nature Chemical Biology*, 16(4):458–468, Apr. 2020. ISSN 1552-4469. doi: 10.1038/s41589-019-0425-0.

A. Popa, K. Lebrigand, A. Paquet, N. Nottet, K. Robbe-Sermesant, R. Waldmann, and P. Barbry. RiboProfiling: A Bioconductor package for standard Ribo-seq pipeline processing [version 1; peer review: 3 approved]. *F1000Research*, 5(1309), 2016. doi: 10.12688/f1000research.8964.1.

A. Raj, S. H. Wang, H. Shim, A. Harpak, Y. I. Li, B. Engelmann, M. Stephens, Y. Gilad, and J. K. Pritchard. Thousands of novel translated open reading frames in humans inferred by ribosome footprint profiling. *eLife*, 5:e13328, May 2016a. ISSN 2050-084X. doi: 10.7554/eLife.13328.

A. Raj, S. H. Wang, H. Shim, A. Harpak, Y. I. Li, B. Engelmann, M. Stephens, Y. Gilad, and J. K. Pritchard. Thousands of novel translated open reading frames in humans inferred by ribosome footprint profiling. *eLife*, 5:e13328, May 2016b. ISSN 2050-084X. doi: 10.7554/eLife.13328.

C. A. Rubio, B. Weisburd, M. Holderfield, C. Arias, E. Fang, J. L. DeRisi, and A. Fanidi. Transcriptome-wide characterization of the eIF4A signature highlights plasticity in translation regulation. *Genome Biology*, 15(10):476, Oct. 2014. ISSN 1474-760X. doi: 10.1186/s13059-014-0476-1.

N. Stern-Ginossar, B. Weisburd, A. Michalski, V. T. K. Le, M. Y. Hein, S.-X. Huang, M. Ma, B. Shen, S.-B. Qian, H. Hengel, M. Mann, N. T. Ingolia, and J. S. Weissman. Decoding Human Cytomegalovirus. *Science*, 338(6110):1088–1093, Nov. 2012. doi: 10.1126/science.1227919.

J. Su, Y. Lu, S. Pan, A. Murtadha, B. Wen, and Y. Liu. RoFormer: Enhanced Transformer with Rotary Position Embedding, Aug. 2022.

M. E. Tanenbaum, N. Stern-Ginossar, J. S. Weissman, and R. D. Vale. Regulation of mRNA translation during mitosis. *eLife*, 4:e07957, Aug. 2015. ISSN 2050-084X. doi: 10.7554/eLife.07957.

A. Werner, S. Iwasaki, C. A. McGourty, S. Medina-Ruiz, N. Teerikorpi, I. Fedrigo, N. T. Ingolia, and M. Rape. Cell-fate determination by ubiquitin-dependent regulation of translation. *Nature*, 525(7570): 523–527, Sept. 2015. ISSN 1476-4687. doi: 10.1038/nature14978.

Z. Xiao, R. Huang, X. Xing, Y. Chen, H. Deng, and X. Yang. De novo annotation and characterization of the translatome with ribosome profiling data. *Nucleic Acids Research*, 46(10):e61, June 2018. ISSN 1362-4962. doi: 10.1093/nar/gky179.

Z. Xu, L. Hu, B. Shi, S. Geng, L. Xu, D. Wang, and Z. J. Lu. Ribosome elongating footprints denoised by wavelet transform comprehensively characterize dynamic cellular translation events. *Nucleic Acids Research*, 46(18):e109, Oct. 2018. ISSN 0305-1048. doi: 10.1093/nar/gky533.

P. Zhang, D. He, Y. Xu, J. Hou, B.-F. Pan, Y. Wang, T. Liu, C. M. Davis, E. A. Ehli, L. Tan, F. Zhou, J. Hu, Y. Yu, X. Chen, T. M. Nguyen, J. M. Rosen, D. H. Hawke, Z. Ji, and Y. Chen. Genome-wide identification and differential analysis of translational initiation. *Nature Communications*, 8(1):1749, Nov. 2017. ISSN 2041-1723. doi: 10.1038/s41467-017-01981-8.

H. Zur, R. Aviner, and T. Tuller. Complementary Post Transcriptional Regulatory Information is Detected by PUNCH-P and Ribosome Profiling. *Scientific Reports*, 6(1):21635, Feb. 2016. ISSN 2045-2322. doi: 10.1038/srep21635.
